# Efficient evolution of human antibodies from general protein language models and sequence information alone

**DOI:** 10.1101/2022.04.10.487811

**Authors:** Brian L. Hie, Duo Xu, Varun R. Shanker, Theodora U.J. Bruun, Payton A. Weidenbacher, Shaogeng Tang, Peter S. Kim

**Affiliations:** Department of Biochemistry, Stanford University School of Medicine, Stanford, CA 94305, USA; Stanford ChEM-H, Stanford, CA 94305, USA; Stanford Medical Scientist Training Program, Stanford University School of Medicine, Stanford CA 94305, USA; Department of Chemistry, Stanford University, Stanford, CA 94305, USA; Chan Zuckerberg Biohub, San Francisco, CA 94158, USA

## Abstract

Natural evolution must explore a vast landscape of possible sequences for desirable yet rare mutations, suggesting that learning from natural evolutionary strategies could accelerate artificial evolution. Here, we report that deep learning algorithms known as protein language models can evolve human antibodies with high efficiency, despite providing the models with no information about the target antigen, binding specificity, or protein structure, and also requiring no additional task-specific finetuning or supervision. We performed language-model-guided affinity maturation of seven diverse antibodies, screening 20 or fewer variants of each antibody across only two rounds of evolution. Our evolutionary campaigns improved the binding affinities of four clinically relevant antibodies up to 7-fold and three unmatured antibodies up to 160-fold across diverse viral antigens, with many designs also demonstrating improved thermostability and viral neutralization activity. Notably, our algorithm requires only a single wildtype sequence and computes recommended amino acid changes in less than a second. Moreover, the same models that improve antibody binding also guide efficient evolution across diverse protein families and selection pressures, indicating that these results generalize to many natural settings. Contrary to prevailing notions of evolution as difficult and resource-intensive, our results suggest that when constrained to a narrow manifold of evolutionary plausibility, evolution can become much easier, which we refer to as the “efficient manifold hypothesis.”

## Introduction

An apparent paradox in evolutionary biology is how a random process can reliably generate new functions in short timescales across settings as diverse as antibody affinity maturation, viral immune escape, or tumor evolution [1]–[6]. The paradox arises from the difficulty of the task (exploring an immense space of possible sequences for rare mutations that improve fitness) contrasted with a seemingly simple set of tools (random mutation and recombination). Current approaches for directed evolution of proteins in the laboratory [7] illustrate this contrast, as high-throughput evolutionary screens that rely on random guessing or brute-force search often devote substantial effort to interrogating weakly active or nonfunctional proteins.

One approach to improve the efficiency of artificial evolution is to learn the rules of evolutionary plausibility (for example, sequences that result in a valid antibody) to help bias evolution away from invalid regimes (for example, mutations that cause an antibody to misfold) [8]. However, even if a search space were restricted to a set of evolutionarily plausible antibodies, the subset of those antibodies with improved binding affinity to a specific target might still be rare beyond practical utility (**Figure 1A**). More broadly, a major open question [9] is whether learning general evolutionary rules, or “intrinsic fitness,” is sufficient to enable efficient evolution under specific definitions of “extrinsic fitness” (for example, high binding affinity).

**Figure 1:**
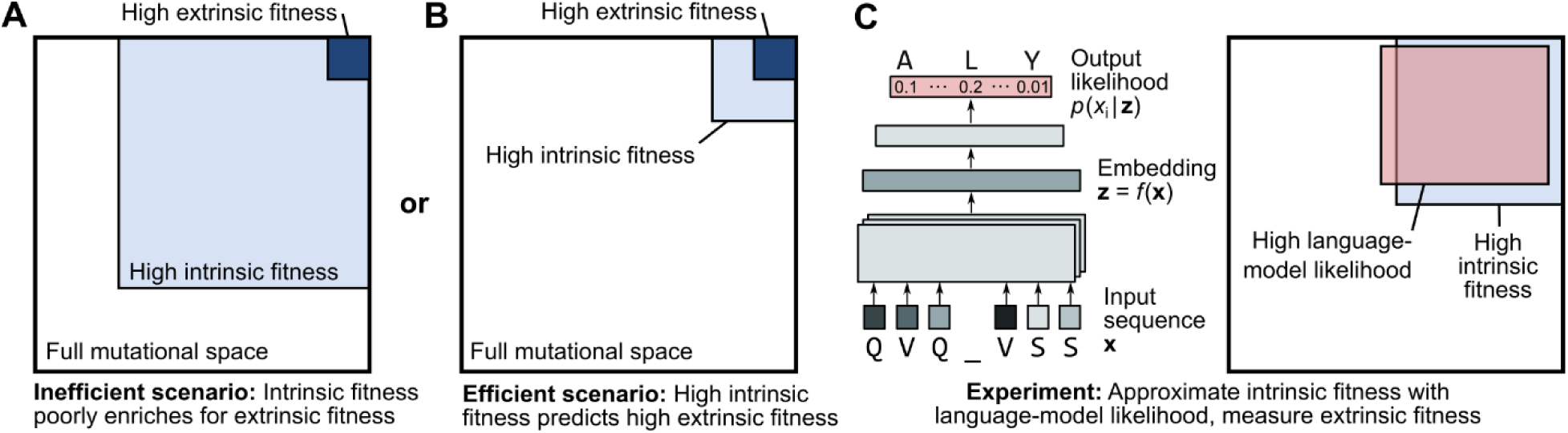
Guiding evolution with protein language models. (**A, B**) Two possible models for relating the space of mutations with high evolutionary plausibility, or intrinsic fitness (for example, the set of valid antibodies), to the space with high fitness under specific selection pressures, or extrinsic fitness (for example, the set of antibodies with high binding affinity to a specific antigen). Both models assume that mutations with high extrinsic fitness make up a rare subset of the full mutational space. Under the first model (**A**), mutations with high extrinsic fitness are also rare within the subset of mutations with high intrinsic fitness. Under the second model (**B**), when restricted to the regime of high intrinsic fitness, mutations with high extrinsic fitness become much more common. (**C**) Protein language models, trained on millions of natural protein sequences, learn amino-acid patterns that are likely to occur in nature. We thus hypothesize that language-model likelihood approximates intrinsic fitness. Assuming that this is a good approximation, and if the second model (**B**) better describes nature, then a language model with no information about specific selection pressures can still efficiently guide evolution.

Here we show that evolutionary information alone can lead to improved fitness under specific selection pressures with high efficiency (**Figure 1B**). For our main experimental test case, we focus on affinity maturation of human antibodies in which high fitness is defined as stronger binding affinity to a particular antigen. In vivo, a process known as somatic hypermutation evolves or “matures” an antibody lineage to have higher affinity for an antigen via repeated mutagenesis [4], [10], [11]. Ex vivo, affinity maturation is a major application of directed evolution due to the therapeutic potential of antibodies with high affinity for disease targets [12].

To model evolutionary plausibility, we use algorithms known as neural language models (**Figure 1C**), which are trained on large datasets of sequences to learn patterns that are likely to occur in natural proteins [13]–[21]. Importantly, we use general language models [17], [18] trained on sequence datasets that are meant to represent variation across all observed natural proteins [22], rather than a language model that is restricted to variation among antibodies [23]–[26]. Given a single starting sequence, we use these language models to recommend plausible amino acid substitutions that we then experimentally screen for improved fitness. We design our approach to be highly general: the algorithm requires only a single wildtype sequence, without any initial binding affinity data, knowledge of the antigen, task-specific supervision, evolutionary homologs, or protein structure information, and can recommend changes to the wildtype sequence in seconds.

With this approach, we evolve seven human immunoglobulin G (IgG) antibodies that bind to antigens from coronavirus, ebolavirus, and influenza A virus. We focus on viral antigens given the importance of antibody therapeutics for epidemic and pandemic viral diseases [27]–[30]. When evolving clinically relevant antibodies, which are already highly mature, our best design achieves a 7-fold improvement in binding affinity from wildtype; for unmatured antibodies, our best design achieves a 160-fold improvement. Many of the designs also preserve or improve thermostability and pseudovirus neutralization activity, including a significant improvement in the neutralization potency of a clinically approved therapeutic antibody for ebolavirus disease (Ebola). We measure 20 or fewer new variants of each antibody across just two rounds of evolution, which, to our knowledge, represents unprecedented efficiency for machine learning-guided directed evolution [31], [32] and which supports the practical utility of our approach. This performance is especially striking given that the underlying language models are completely unsupervised and have no initial task-specific training data. Also notable is that around half of the amino acid substitutions that improve affinity are located in antibody framework regions, which are much less mutated during natural affinity maturation [11] and are thus often excluded from artificial evolution [33], [34].

Beyond antibodies, we show that this efficiency applies generally across other protein families as well. In particular, we demonstrate that the *same* general protein language models that we used to affinity-mature antibodies can also predict antibiotic resistance, enzyme activity, or viral replication fitness. Our results suggest that evolution guided by language models offers a compelling alternative to brute-force search, random guessing, or even rational design as a strategy for evolving proteins in the laboratory. Moreover, we discuss how the success of our approach challenges existing notions of evolutionary difficulty by suggesting that natural evolutionary manifolds (that is, the set of intrinsically plausible sequences) are efficiently primed for fitness-enhancing mutations under extrinsic selection pressures.

## Results

### Efficient affinity maturation with general protein language models

Recent work has demonstrated that language models can predict natural evolution despite having no knowledge of specific selection pressures [9]. However, this prior work only predicted the direction of evolution retrospectively when given full knowledge of the evolutionary trajectory. We therefore sought to investigate if the same language models could predict unobserved evolution to prospectively design new proteins.

In particular, we hypothesized that the predictive capabilities of protein language models might enable a researcher to provide only a single, wildtype antibody sequence to the algorithm and receive a small, manageable set (∼10^1^) of high-likelihood variants to experimentally measure for desirable properties. This is a very general setting that does not assume knowledge of protein structure or task-specific training data, thereby avoiding the resource-intensive processes associated with structure determination [34] or high-throughput screens [33]. A major question, however, is if higher evolutionary likelihood would efficiently translate to higher fitness.

We tested our hypothesis by conducting separate directed evolution campaigns, guided by language-model likelihood, to affinity-mature seven antibodies representing diverse antigens and degrees of maturity (**Supplementary Table 1**). These antibodies are:

- MEDI8852: A broadly neutralizing antibody (bnAb) that binds influenza A hemagglutinin (HA) across variants of both major phylogenetic groups (Group 1 and Group 2) and that reached Phase-II clinical trials; this antibody is highly matured, with its parent being isolated from a human followed by substantial artificial evolution [27].
- MEDI8852 unmutated common ancestor (UCA): The unmatured, inferred germline sequence of MEDI8852, which only neutralizes viruses with Group 1 HAs [27].
- mAb114: A patient-derived antibody that neutralizes ebolavirus by binding to its glycoprotein (GP) [28] and has been approved for clinical use by the United States Food and Drug Administration (FDA).
- mAb114 UCA: The unmatured, inferred germline sequence of mAb114 with weak binding to ebolavirus GP [28].
- S309: A patient-derived antibody that cross-neutralizes the sarbecoviruses SARS-CoV-1 (severe acute respiratory syndrome coronavirus 1) and SARS-CoV-2 by binding to the spike glycoprotein (Spike) [29] and is the parent antibody of sotrovimab [35], which currently has FDA emergency-use authorization (EUA) for treatment of COVID-19 (coronavirus disease 2019).
- REGN10987: A patient-derived antibody that binds early variants of SARS-CoV-2 Spike [30] and that had an FDA EUA for use against these variants.
- C143: An unmatured, patient-derived antibody that binds the SARS-CoV-2 Wuhan-Hu-1 Spike but was isolated prior to extensive in-vivo somatic hypermutation [36], [37].

We performed evolution with the ESM-1b language model and the ESM-1v ensemble of five language models (six language models in total) [17], [18]. ESM-1b and ESM-1v were trained on UniRef50 and UniRef90, respectively, which are protein sequence datasets that represent variation across all observed natural proteins and include only a limited number of antibody sequences (UniRef90 contains ∼800 immunoglobulin variable regions out of ∼98 million total sequences) [22]. These datasets are also constructed such that no two sequences have more than 50% (UniRef50) or 90% (UniRef90) sequence similarity with each other, and immunoglobulin sequences are further curated to avoid biological redundancy. Additionally, because both datasets precede the COVID-19 pandemic, any SARS-CoV-2-related antibody sequence variation would not be included. Therefore, to evolve these antibodies, the language models cannot leverage antigen- or disease-specific biases in the training data and must instead learn more intrinsic evolutionary patterns.

We used these language models to compute likelihoods of all single-residue substitutions to the antibody variable regions of either the heavy chain (VH) or the light chain (VL). We selected substitutions with higher evolutionary likelihood than wildtype across a consensus of six language models; additional details are provided in **Methods**. In the first round of evolution, we measured the antigen-binding affinity by biolayer interferometry (BLI) of variants that only contain a single-residue substitution from wildtype. In the second round, we measured variants containing combinations of substitutions, where we selected substitutions that corresponded to preserved or improved binding based on the results of the first round. We performed these two rounds for all seven antibodies, measuring 8 to 14 variants per antibody in round one and 1 to 11 variants per antibody in round two (**Figure 2**, **Supplementary Table 1**). Variants of the clinically relevant antibodies, which have very low or undetectable dissociation as IgGs, were screened by measuring the dissociation constant (*K*_d_) of the monovalent fragment antigen-binding (Fab) region; variants of the unmatured antibodies were screened by measuring the apparent *K*_d_ of the bivalent IgG, followed by also measuring the *K*_d_ values of the Fab fragments of the highest-avidity variants (**Methods**).

**Figure 2:**
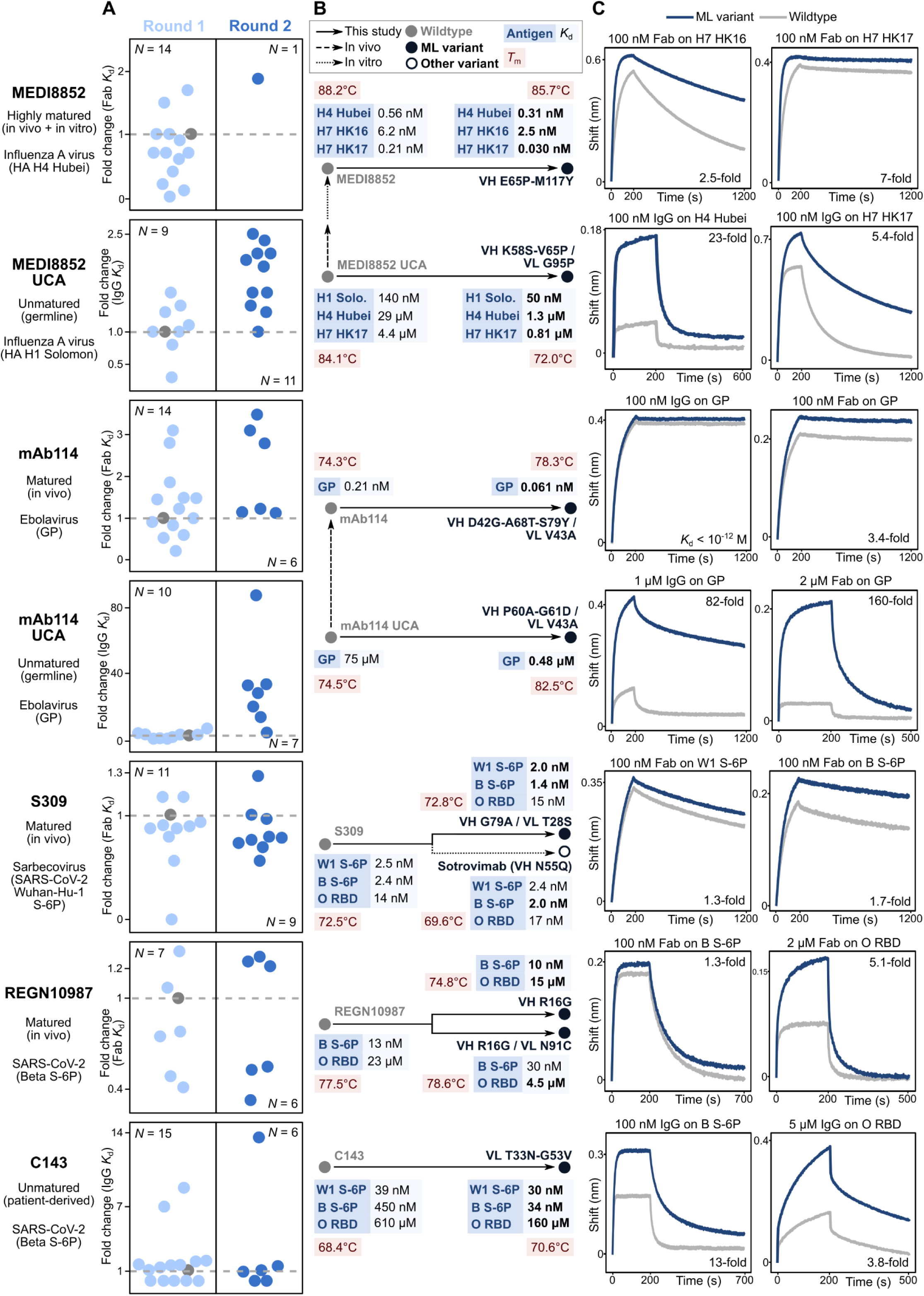
Language-model-guided affinity maturation of seven human antibodies. (**A**) Strip plots visualizing the two rounds of directed evolution conducted for each antibody. Each point represents an IgG or Fab variant plotted according to the fold-change in *K*_d_ from wildtype on the *y*-axis and jitter on the *x*-axis; a gray, dashed line is drawn at a fold change of 1 and the wildtype point is colored gray. MEDI8852 variants were screened against HA H4 Hubei, MEDI8852 UCA variants against HA H1 Solomon, mAb114 and mAb114 UCA variants against ebolavirus GP, S309 variants against Wuhan-Hu-1 S-6P, and REGN10987 and C143 variants against Beta S-6P. (**B**) Phylogenetic trees illustrating the evolutionary trajectories from wildtype to the highest-affinity variant(s) of each antibody. Nodes are annotated with the *K*_d_ values for different antigens and the *T*_m_ of the Fab; all *K*_d_ values are for the monovalent Fab versions except those of C143, which are apparent *K*_d_ values for the bivalent IgGs. Bolded *K*_d_ values indicate a 1.1-fold or higher improvement from wildtype. ML variant: machine-learning-guided variant; H1 Solo.: H1 Solomon; W1: Wuhan-Hu-1; B: Beta, O: Omicron. (**C**) We obtained avidity and affinity measurements via biolayer interferometry (BLI) of IgGs and Fabs at the indicated concentrations binding to the indicated antigen. Selected BLI traces of the highest-affinity variants for the respective antigens are plotted alongside those of the wildtype variants.

We could successfully express all but one of 122 new variants across our seven evolutionary trajectories, indicating that language-model likelihood is a good approximation of evolutionary plausibility (**Figure 1C**). Across all seven antibodies, we found that 71% to 100% of the first-round Fab variants (containing a single-residue substitution) retained sub-micromolar binding to the antigen, and 15% to 71% percent of first-round variants led to improved binding affinity (defined as a 1.1-fold or higher improvement in *K*_d_ compared to wildtype) (**Supplementary Table 1**). Most of the second-round variants (containing a combination of substitutions) also have improved binding, with some combinations demonstrating additive or synergistic effects (**Supplementary Tables 1-9**). 36 out of all 76 language-model-recommended, single-residue substitutions (and 18 out of 32 substitutions that lead to improved affinity) occur in framework regions (**Supplementary Tables 2-9**), which are generally much less mutated during conventional affinity maturation compared to the complementarity-determining regions (CDRs) [11].

We were able to improve the binding affinities for all clinically relevant antibodies tested, despite these antibodies being already highly evolved (starting at low nanomolar or picomolar affinity). MEDI8852 is a potent binder with a sub-picomolar Fab *K*_d_ across many HAs and picomolar or nanomolar binding to HAs from subtypes H4 and H7. While we explicitly screened variants using an HA H4 antigen, the best design also improves binding across a broad set of HAs (**Supplementary Tables 2** and **3**), including a 7-fold improvement (from 0.21 nM to 0.03 nM) for HA H7 HK17 (A/Hong Kong/125/2017(H7N9)). The best variant of mAb114, a clinically approved drug, achieves a 3.4-fold improvement in Fab *K*_d_ for ebolavirus GP (**Supplementary Table 5**). For REGN10987, the highest-affinity variant has a 1.3-fold improvement against Beta-variant Spike with six stabilizing proline substitutions (S-6P) [38] (the antigen used in screening), while another of our designs has a 5.1-fold improvement for the Omicron-variant receptor binding domain (RBD) (**Supplementary Table 8**). For S309, we compared our designs to wildtype and to a variant with the N55Q substitution in the VH introduced after a small-scale, rational evolutionary screen [35]; the S309 Fab with the VH N55Q substitution forms the Fab of the therapeutic antibody, sotrovimab. Interestingly, our best variant of S309 has higher affinity than sotrovimab, including a 1.3-fold improvement in Fab *K*_d_ compared to wildtype S309 (versus 1.1-fold for sotrovimab) for SARS-CoV-2 Wuhan-Hu-1 S-6P (the antigen used in screening), a 1.7-fold improvement (versus 1.3-fold for sotrovimab) for Beta S-6P, and a 0.93-fold change (versus 0.82-fold for sotrovimab) for Omicron RBD (**Supplementary Table 7**).

We were also able to improve affinities for all three unmatured antibodies, often involving much higher fold changes than when evolving the matured antibodies, indicating easier evolvability with respect to affinity. For MEDI8852 UCA, the best Fab design achieves a 2.6-fold improvement in *K*_d_ against HA H1 Solomon (A/Solomon Islands/3/2006(H1N1)), the antigen used in screening. Our best designs also acquire breadth of binding to some Group 2 HAs, including a 23-fold improvement for HA H4 Hubei (A/swine/Hubei/06/2009(H4N1)) and a 5.4-fold improvement for HA H7 HK17 (**Supplementary Table 4**). For mAb114 UCA, our best Fab design achieves a 160-fold improvement in *K*_d_ for ebolavirus GP (**Supplementary Table 6**). Strikingly, despite having no knowledge about the matured form, the algorithm recommends amino acid substitutions to both of these UCA antibodies that are also observed in the matured antibody. Interestingly, other affinity-enhancing substitutions to the UCA antibodies are not found in the matured versions: excluding any substitutions or modified sites found in the matured antibody, our UCA variants achieve up to a 7-fold improvement for HA H4 Hubei (variant VH P75R / VL G95P; **Supplementary Table 4**) and a 33-fold improvement for ebolavirus GP (variant VH G88E / VL V43A; **Supplementary Table 6**), demonstrating that our algorithm successfully explores alternative evolutionary routes. For C143, a patient-derived antibody isolated prior to extensive affinity maturation [37], our best design achieves a 13-fold improvement for Beta S-6P and a 3.8-fold improvement for Omicron RBD (**Supplementary Table 9**). Results from our directed evolution campaigns are further summarized in **Figure 2**, **Supplementary Tables 2-9**, and **Supplementary Data 1**. In total, across antibodies representing diverse antigens and degrees of maturity, our approach consistently and efficiently produces higher-affinity variants.

### Improved thermostability and neutralization of evolved antibodies

Although we explicitly selected for variants with improved binding to specific antigens, we also sought to establish if these variants have improved stability (**Methods**). We found that Fabs for 21 out of the 31 language-model-recommended, affinity-enhancing variants that we tested had a higher melting temperature (*T*_m_) than wildtype, and all variants maintained thermostability (*T*_m_ > 70°C). When evolving S309 to have higher affinity, our best design has a *T*_m_ of 72.8°C compared to 72.5°C for wildtype, whereas the VH N55Q substitution introduced in sotrovimab decreases the *T*_m_ to 69.6°C (**Figure 2**). Our evolved variants for mAb114, mAb114 UCA, REGN10987, and C143 also preserve or improve *T*_m_; the highest change we observed was an increase from 74.5°C to 82.5°C when evolving mAb114 UCA. Improved thermostability does not completely explain our affinity maturation results, however, as we observe somewhat decreased *T*_m_ for our affinity-matured variants of MEDI8852 and its UCA, though these Fabs are still thermostable (**Figure 2**).

We also wanted to determine if our affinity-matured variants have better viral neutralization activity. We tested affinity-enhancing variants of four antibodies using pseudovirus neutralization assays (**Methods**) and in all cases observed variants with half-maximal inhibitory concentration (IC_50_) values that are significantly improved (Bonferroni-corrected, one-sided *t*-test *P* < 0.05), including a 1.5-fold improvement for the best mAb114 variant against Ebola pseudovirus, a 2-fold improvement for the best REGN10987 variant against SARS-CoV-2 Beta pseudovirus, and a 32-fold improvement for the best C143 variant against Beta pseudovirus (**Figure 3A** and **Supplementary Fig. 1**; **Supplementary Tables 5, 8,** and **9**). Additionally, while the IC_50_ values of variants of mAb114 UCA are greater than the highest tested concentration, the affinity-matured variants demonstrate detectable neutralization at a >100-fold lower concentration compared to wildtype (**Supplementary Fig. 1**). In general, change in binding affinity corelates well with change in neutralization (Spearman *r* = 0.82, two-sided *t*-distribution *P* = 1.9 × 10^-4^) (**Figure 3B**). Given the limited number of variants tested, we also note that alternative versions of our directed evolution campaigns could have instead explicitly screened variants for neutralization activity.

**Figure 3:**
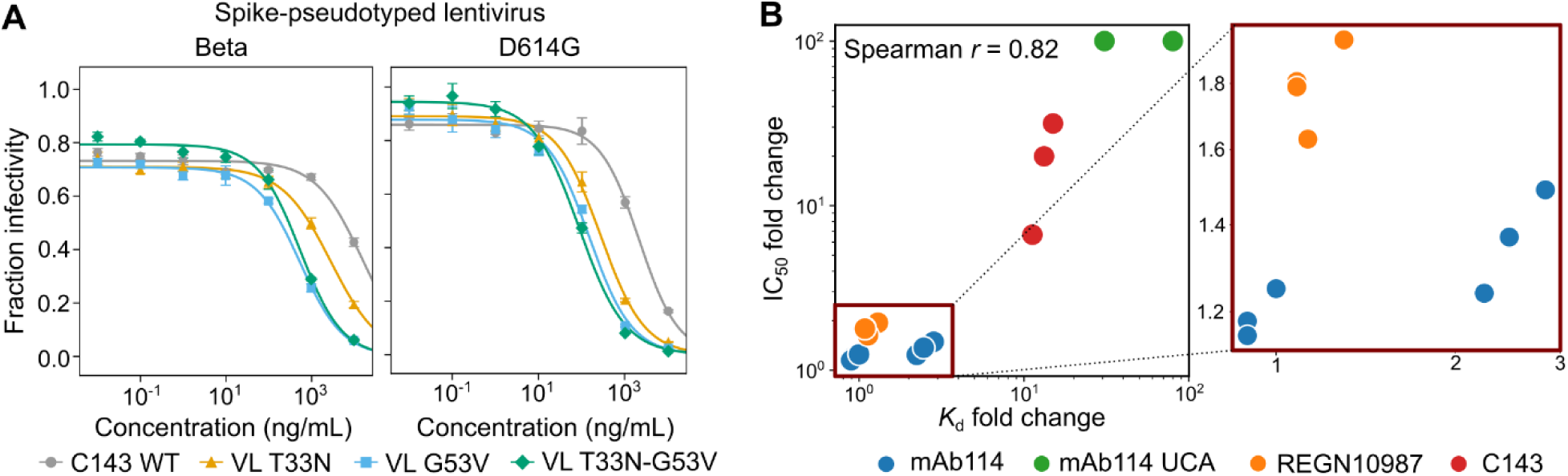
Affinity-matured variants improve pseudovirus neutralization. (**A**) Variants of the antibody, C143, obtained from our language-model-guided affinity maturation campaign demonstrate improved neutralization activity in a pseudovirus assay. For Beta pseudovirus, out of the three higher-affinity variants that we also screened for neutralization activity, the best improvement is the 32-fold improvement of VL G53V; for D614G pseudovirus, the best improvement is the 19-fold improvement of VL T33N-G53V (**Supplementary Table 9**). Also see **Supplementary** Fig. 1. Points indicate the mean; error bars indicate the standard deviation. (**B**) Fold-change in *K*_d_ correlates well with fold-change in IC_50_ (Spearman *r* = 0.82) across all designs tested, consistent with higher binding affinity contributing to improved viral neutralization activity.

### Originality of affinity-enhancing substitutions

While the ability to find any improvement in affinity is itself useful for engineering applications, we were also interested in whether some of the changes recommended by our algorithm demonstrate “originality.” We quantified originality by computing the frequency that a given residue is observed in nature (**Methods**), where a change to a rarely observed residue indicates that the model learns patterns that go beyond its literal training dataset. While many affinity-enhancing substitutions are indeed observed at high frequency in both the model’s training data [22] and in a database of antibody sequences [39], other substitutions demonstrate greater originality. For example, in the MEDI8852 UCA trajectory, the VL G95P framework substitution (**Figure 2**; **Supplementary Table 4**) involves changing a glycine observed in 99% of natural antibody sequences to a proline observed in <1% of natural sequences. Overall, five out of 32 affinity-enhancing substitutions (∼16%) involve changing the wildtype residue to a rare or uncommon residue (**Supplementary Table 10**). These results indicate that the language models learn both the “easy” evolutionary rules involving high-frequency residues and more complex rules that are not captured by a multiple sequence alignment or conventional antibody evolution. Conceptually, these low-frequency, affinity-enhancing substitutions are analogous to examples in other disciplines where an artificial-intelligence program occasionally makes unusual but advantageous choices (for example, unintuitive game-playing decisions [40]), and likewise may be worth further study.

### Generality across diverse protein families

Given the success of general protein language models at guiding antibody evolution, we also tested how well the same models could acquire high-fitness variants across a range of protein families. Previous work has demonstrated that the likelihoods from general protein language models have good correlation with experimental phenotypes from high-throughput assays over ∼10^3^ to 10^4^ variants [9], [18]. Previous computational simulations have also indicated that these models can help bias multi-round evolution away from large regions of a sequence landscape with zero or very low fitness [8].

Here, we observe that the same models we used to affinity-mature antibodies can also guide efficient evolution when measuring only a small number (∼10^1^) of variants according to diverse definitions of extrinsic fitness including antibiotic resistance, cancer drug resistance, enzyme activity, or viral replication fitness [41]. More specifically, we used the same algorithm and language models in our affinity-maturation experiments to instead suggest changes to wildtype sequences from human, bacterial, or viral organisms representing eight diverse protein families. We then used fitness measurements from high-throughput scanning mutagenesis experiments [41], [42] to validate the language-model-recommended predictions (notably, these measurements were not provided to the model).

Across diverse proteins, language-model-recommended variants are significantly enriched (hypergeometric *P* < 0.05) for high fitness values, and high-fitness variants make up a much larger portion of language-model-recommended variants compared to random guessing in nearly all cases (**Figure 4A**, **Supplementary Fig. 2**, and **Supplementary Table 11**). For example, while ampicillin resistance is observed for just 7% of all single-residue substitutions to β-lactamase, it is observed for 40% of language-model-recommended substitutions, and the same set of language models can also help prioritize single-residue substitutions to HA that result in high viral infectivity (from 7% to 31%) and substitutions to PafA that improve enzyme kinetics (from 3% to 20%). Additionally, across all proteins, even the first round of a small-scale evolutionary campaign guided by language models would yield variants that are near the local fitness peak (**Supplementary Fig. 2**). In total, these results suggest that the evolutionary efficiency that we observed for affinity-maturation of human IgGs also generalizes to diverse natural settings.

**Figure 4:**
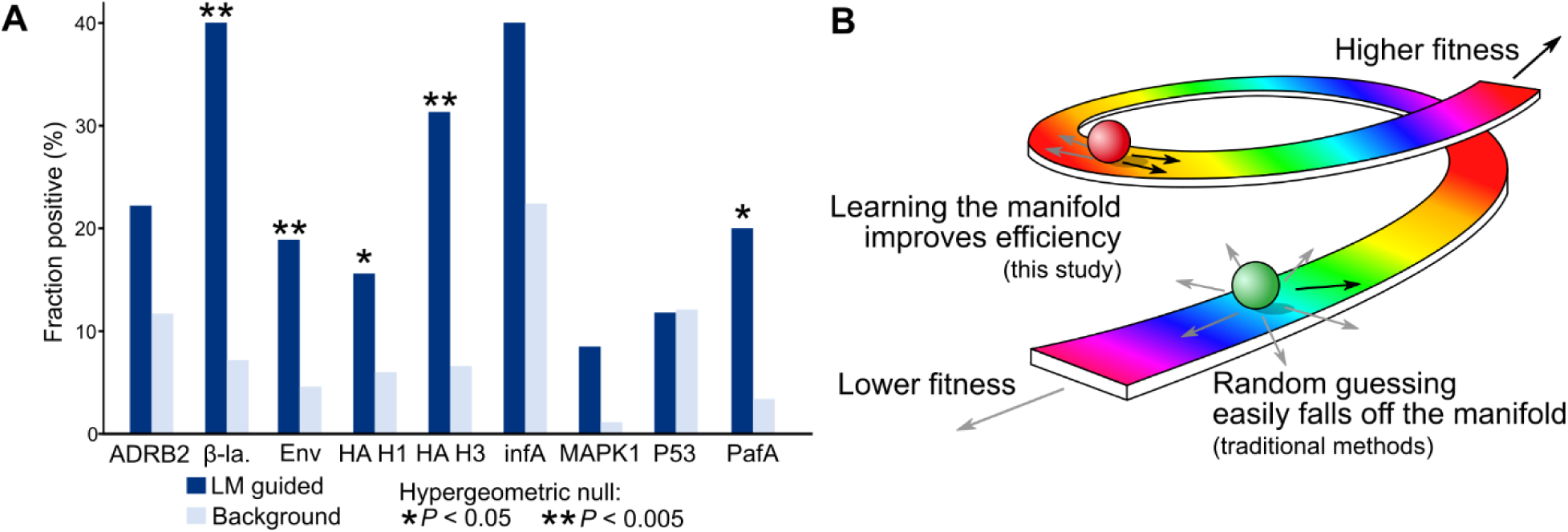
The efficient manifold hypothesis. (**A**) The *same* strategy and language models that we use to affinity-mature antibodies can also recommend high-fitness changes across a diversity of protein families selection pressures, as identified experimentally using high-throughput scanning mutagenesis assays [41], [42] (described in **Supplementary Table 11**). “Fraction positive” indicates the percentage of high-fitness amino acid substitutions within either the set of substitutions recommended by the language model (LM guided) or the set of all single-residue substitutions (Background). A substantial portion (12% to 40%) of language-model-guided substitutions have high extrinsic fitness, which in many cases is significantly enriched compared to the background percentage; also see **Supplementary** Fig. 2 and **Supplementary Table 11**. ADRB2: adrenoreceptor beta 2; β-la.: β-lactamase; Env: envelope glycoprotein; infA: translation initiation factor 1; MAPK1: mitogen-activated protein kinase 1; PafA: phosphate-irrepressible alkaline phosphatase. (**B**) Conceptually, intrinsic fitness forms a manifold that is represented in this cartoon by the rainbow road, where ascending corresponds to improving extrinsic fitness and descending corresponds to lowering extrinsic fitness. Under the efficient manifold hypothesis, this manifold of intrinsic fitness is narrow, therefore moving in any direction (for example, via random or brute-force mutagenesis) would most likely decrease extrinsic fitness or fall off the manifold entirely (represented by the green ball). However, if movement is constrained to the narrow manifold of intrinsic fitness (for example, when guided by a language model), then the chance of improving extrinsic fitness increases substantially (represented by the red ball).

## Discussion

We show that general protein language models can guide highly efficient affinity maturation based on the wildtype antibody sequence alone. We improved binding affinities of a highly evolved influenza A broadly neutralizing antibody (bnAb), MEDI8852, by up to 7-fold and a clinically approved ebolavirus antibody, mAb114, by 3.4-fold. We also evolve S309, a sarbecovirus bnAb, to have higher affinity and thermostability than a rationally designed and clinically available variant, sotrovimab. We improved binding affinities of unmatured antibodies from 13- to 160-fold across diverse antigens, which is within the 3.8- to 580-fold improvement range previously achieved by a state-of-the-art, in-vitro evolutionary system applied to unmatured, anti-RBD nanobodies (in which the computational portion of our approach, which takes seconds, is replaced with rounds of cell culture and sorting, which takes weeks) [12]. We also note that in-vitro, cell-surface-display methods encounter physical limits that make it challenging to distinguish better binders when the wildtype binder already has high affinity (<1 nM) [43], which is not a limitation of our approach. Moreover, our algorithm is based on language models (trained on general protein sequence variation) that can also predict high-fitness variants across diverse protein families and engineering applications. We envision our approach as useful within preclinical development as a rapid way to identify improved variants of an existing protein of interest (for example, an antibody isolated from a patient or from a naïve library). We also anticipate that language models will become a key part of the antibody engineer’s toolkit.

Interestingly, about half of the language-model-recommended substitutions (and about half of the affinity-enhancing substitutions) fall in framework regions, which are typically not proximal to the binding interface and are therefore sometimes excluded from directed evolution [33]. While some of these framework changes may improve affinity via protein stabilization, others do not appear to increase thermostability and may instead be causing larger-scale rearrangements that improve affinity via structural reorientation, which has been observed in natural affinity maturation [44]–[46]. Our algorithm also recommends a number of affinity-enhancing substitutions with low observed frequency in nature. An interesting area for future work is to characterize additional biochemical or structural features of these unconventional changes.

The broader relevance of our results, beyond affinity maturation of human antibodies, arises from asking why this method works. Fundamentally, our results are surprising in that modifying amino acid residues simply based on evolutionary plausibility, or “intrinsic fitness,” sufficiently enriches for changes that improve fitness under specific, natural selection pressures, or “extrinsic fitness” (**Figure 1B**). These results challenge a prevailing notion that evolution is difficult because it is random. Instead, we hypothesize that, in many settings, as long as evolution remains on a naturally plausible manifold, a substantial portion (greater than 10%) of mutations are bound to improve extrinsic fitness, which we call the “efficient manifold hypothesis” (**Figure 4B**). Our findings for both antibodies and other natural proteins provide direct support for this hypothesis. The efficient manifold hypothesis is also supported by the recent successes of completely unsupervised models in predicting evolution under a variety of specific selection pressures, from clinical variant risk to viral fitness and immune escape potential [15], [47]–[49].

The efficient manifold hypothesis has direct, practical applications for those trying to evolve proteins in the laboratory. Evolution guided by a language model can be used as a drop-in replacement for current evolutionary tools based on randomization; for example, combinatorial libraries [50], [51] can recombine language-model-guided mutations alongside or instead of rationally chosen mutations [33]. By leveraging increasingly efficient technologies for nucleic acid printing [42], language-model-guided evolution could also directly replace mutagenesis strategies based on, for example, an error-prone polymerase.

The efficient manifold hypothesis also challenges the notion that explicitly modeling extrinsic fitness with a supervised model should be the best or the default approach to machine-learning-guided directed evolution [32]. To the end user, guiding evolution via pretrained, unsupervised models is less resource-intensive than collecting enough task-specific data to train a supervised model [33]. Our techniques can also be used in conjunction with supervised approaches [8], [31]–[34], [52]–[55], and supervising a model over multiple experimental rounds might ultimately lead to higher fitness. However, in many practical settings (for example, the rapid development of sotrovimab in response to the COVID-19 pandemic [35]), the efficiency of an unsupervised, single-round approach is preferable to a protracted, multi-round (machine-learning-guided) directed evolution campaign.

We note that taking advantage of the efficient manifold hypothesis to improve extrinsic fitness may be more difficult when the selection pressure is unnatural or if the wildtype sequence is already at a fitness peak. Relatedly, a potential limitation of our specific algorithm is that we use language models that are trained only on natural sequences and might therefore be less applicable to unnatural proteins generated via de-novo design [56], [57]. However, in many practical design tasks, natural sequences and selection pressures are already preferrable; for example, therapeutic development often prefers human antibodies due to considerations of immunogenicity and toxicity.

Beyond protein engineering applications, the efficient manifold hypothesis may also provide new insight into natural evolution. Our results suggest that many natural evolutionary processes occur on efficient manifolds, which may explain how some proteins are able to quickly and consistently acquire new functions; for example, human immunodeficiency virus relies on rapid and substantial intra-host evolution (on the scale of hours to days) to accomplish both infection and transmission [3]. Nature could support efficient manifolds via a number of mechanisms: for example, a recent analysis of *Arabidopsis thaliana* evolution suggests that epigenomic features enable mutations to be intrinsically biased away from implausible choices [42]. If epigenomic or other mechanisms predispose mutations to have high intrinsic fitness, then natural evolution on an efficient manifold would also become easy.

## Methods

### Acquiring amino acid substitutions via language model consensus

We aim to acquire variants for experimental measurement with high predicted evolutionary plausibility and therefore select amino acid substitutions recommended by a consensus of language models. We take as input a single wildtype sequence **x** = (*x*_1_, … , *x_N_*) ∈ 𝒳*^N^*, where 𝒳 is the set of amino acids and *N* is the sequence length. We also require a set of masked language models, which are pretrained to produce conditional likelihoods 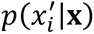. To guide evolution based on a certain language model, we first compute the set substitutions with higher language-model likelihood than the wildtype, i.e., we compute the set

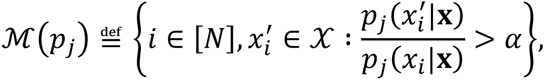

where *p_j_* denotes the language model, *x_i_* denotes the wildtype residue, and *α* = 1. To further filter substitutions to only those with the highest likelihood, we choose substitutions based on a consensus scheme, where, for a new amino acid 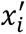, we compute

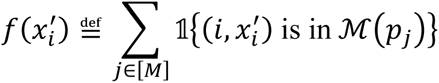

where 𝟙{·} denotes the indicator function and there are *M* language models. We then acquire the set of substitutions with higher likelihood than wildtype across multiple language models, i.e., we acquire

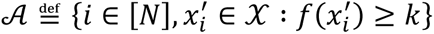

where *k* is a user-supplied cutoff that controls the number of corresponding variants to measure. While we focus on values of *k* that result in small values of |𝒜| (around 10) that can be screened via low-throughput assays, the number of substitutions can be increased by reducing the value of *k* or by lowering the cutoff stringency *α*.

We used six large-scale masked language models, namely, the ESM-1b model [17] and the five models that are ensembled together to form ESM-1v [18], both obtained from https://github.com/facebookresearch/esm. ESM-1b was trained on the March 2018 release of UniRef50 [22] consisting of ∼27 million sequences, and the five models in ESM-1v were each trained on the March 2020 release of UniRef90 [22] consisting of ∼98 million sequences.

### Antibody sequence analysis and evolution

For antibodies, we performed the above steps for the VH and VL sequences separately, obtaining respective sets 𝒜_VH_ and 𝒜_VL_. For round 1 of evolution, we set *α* = 1 and chose values of *k* such that |𝒜_VH_ ∪ *A*_VL_| is approximately 10, which is meant to be a reasonable number of antibody variants for one person to express and purify in parallel. We used *k* = 2 for MEDI8852 VH and VL, *k* = 2 for MEDI8852 UCA VH and VL, *k* = 4 for mAb114 VH and VL, *k* = 2 for mAb114 UCA VH and VL, *k* = 2 for S309 VH, *k* = 1 for S309 VL, *k* = 2 for REGN10987 VH and VL, and *k* = 2 for C143 VH and VL. We further reduced the size of |𝒜_VH_ ∪ *A*_VL_| by requiring the substitution to have the highest likelihood at its respective site for at least one language model. For round 2 of evolution, variants were first measured for binding affinity to a given antigen via BLI (more details below) and those that preserved or enhanced affinity were recombined such that the second-round variants have two or more substitutions from wildtype. For MEDI8852 and MEDI8852 UCA, we tested all possible combinations; for the other antibodies, where the number of possible combinations far exceeds ∼10 variants, we manually selected a set of combinations meant to prioritize inclusion of substitutions that resulted in the largest improvements in affinity during the first round.

We used the wildtype sequences provided by the original study authors describing the respective antibodies [27]–[30], [36]. Wildtype VH and VL sequences are provided in **Supplementary Information**. We used the Kabat region definition provided by the abYsis webtool version 3.4.1 (http://www.abysis.org/abysis/index.html) [39] to annotate the framework regions and CDRs within the VH and VL sequences.

### Antibody cloning

We cloned the antibody sequences into the CMV/R plasmid backbone for expression under a CMV promoter. The heavy chain or light chain sequence was cloned between the CMV promoter and the bGH poly(A) signal sequence of the CMV/R plasmid to facilitate improved protein expression. Variable regions were cloned into the human IgG1 backbone; REGN10987 and C143 variants were cloned with a lambda light chain, while variants of all other antibodies were cloned with a kappa light chain. The vector for both heavy and light chain sequences also contained the HVM06_Mouse (P01750) Ig heavy chain V region 102 signal peptide (MGWSCIILFLVATATGVHS) to allow for protein secretion and purification from the supernatant. VH and VL segments were ordered as gene blocks from Integrated DNA Technologies and were cloned into linearized CMV/R backbones with 5X In-Fusion HD enzyme premix (Takara Bio).

### Antigen cloning

HA, GP, Spike, and RBD sequences were cloned into a pADD2 vector between the rBeta-globin intron and β-globin poly(A). HA constructs contain a Foldon trimerization domain. GP and Spike constructs contain a GCN4 trimerization domain. All HAs, GP, Wuhan-Hu-1 S-6P, and Omicron RBD constructs contain an AviTag. All constructs contain a C-terminal 6xHis tag. We used HA sequences from the following strains: A/New Caledonia/20/1999(H1N1) (H1 Caledonia), A/Solomon Islands/3/2006(H1N1) (H1 Solomon), A/Japan/305/1957(H2N2) (H2 Japan), A/Panama/2007/1999(H3N2) (H3 Panama), A/Victoria/3/1975(H3N2) (H3 Victoria), A/swine/Hubei/06/2009(H4N1) (H4 Hubei), A/Vietnam/1203/2004(H5N1) (H5 Vietnam), A/Hong Kong/61/2016(H7N9) (H7 HK16), and A/Hong Kong/125/2017(H7N9) (H7 HK17). We used Ebola GP ectodomain (Mayinga, Zaire, 1976, GenBank: AAG40168.1) with the mucin-like domain deleted (Δ309-489). Spike or RBD sequences were based off wildtype Wuhan-Hu-1 (GenBank: BCN86353.1), Beta (GenBank: QUT64557.1), or Omicron (GenBank: UFO69279.1).

### DNA preparation

Plasmids were transformed into Stellar competent cells (Takara Bio), and transformed cells were grown at 37°C. Colonies were sequence confirmed and then maxi-prepped per the manufacturer’s recommendations (NucleoBond Xtra Maxi; Macherey-Nagel). Plasmids were sterile filtered using a 0.22-μm syringe filter and stored at 4°C.

### Protein expression

All proteins were expressed in Expi293F cells. For proteins containing a biotinylation tag (AviTag), Expi293F cells containing a stable BirA enzyme insertion were used, resulting in spontaneous biotinylation during protein expression. Expi293F cells were cultured in media containing 66% Freestyle/33% Expi media (ThermoFisher) and grown in TriForest polycarbonate shaking flasks at 37°C in 8% carbon dioxide (CO_2_). The day before transfection cells were spun down and resuspended to a density of 3 × 10^6^ cells/mL in fresh media. The following day, cells were diluted and transfected at a density of approximately 3-4 × 10^6^ cells/mL. Transfection mixtures were made by adding the following components: maxi-prepped DNA, culture media, and FectoPro (Polyplus) would be added to cells to a ratio of 0.5μg:100μL:1.3μL:900μL. For example, for a 100 mL transfection, 50 μg of DNA would be added to 10 mL of culture media, followed by the addition of 130 μL of FectoPro. For antibodies, we divided the transfection DNA equally among heavy- and light-chains; in the previous example, 25 μg of heavy-chain DNA and 25 μg of light-chain DNA would be added to 10 mL of culture media. Following mixing and a 10-minute incubation, the example transfection cocktail would be added to 90 mL of cells. The cells were harvested 3-5 days post-transfection by spinning the cultures at >7,000X g for 15 minutes. Supernatants were filtered using a 0.45-μm filter.

### Antibody purification

We purified antibodies using a 5 mL MAb Select Sure PRISM™ column on the ÄKTA pure fast protein liquid chromatography (FPLC) instrument (Cytiva). The ÄKTA system was equilibrated with line A1 in 1X phosphate-buffered saline (PBS), line A2 in 100 mM glycine pH 2.8, line B1 in 0.5 M sodium hydroxide, Buffer line in 1X PBS, and Sample lines in water. The protocol washes the column with A1, followed by loading of the sample in the Sample line until air is detected in the air sensor of the sample pumps, followed by 5 column-volume washes with A1, elution of the sample by flowing of 20 mL of A2 directly into a 50 mL conical containing 2 mL of 1 M tris(hydroxymethyl)aminomethane (Tris) pH 8.0, followed by 5 column volumes of A1, B1, and A1. We concentrated the eluted samples using 50 or 100 kDa cutoff centrifugal concentrators followed by buffer exchange using a PD-10 column (SEPHADEX) that had been preequilibrated into 1X PBS. Purified antibodies were stored at -20°C.

### Antigen purification

All antigens were His-tagged and purified using HisPur™ Ni-NTA resin (ThermoFisher). Cell supernatants were diluted with 1/3rd volume wash buffer [20 mM imidazole, 20 mM 4-(2-hydroxyethyl)-1-piperazineethanesulfonic acid (HEPES) pH 7.4, 150 mM sodium chloride (NaCl) or 20 mM imidazole, 1X PBS] and the Ni-NTA resin was added to diluted cell supernatants. For all antigens except SARS-CoV-2 Spike, the samples were then incubated at 4°C while stirring overnight. SARS-CoV-2 Spike antigens were incubated at room temperature. Resin/supernatant mixtures were added to chromatography columns for gravity-flow purification. The resin in the column was washed with wash buffer (20 mM imidazole, 20 mM HEPES pH 7.4, 150 mM NaCl or 20 mM imidazole, 1X PBS) and the proteins were eluted with either 250 mM imidazole, 20 mM HEPES pH 7.4, 105mM NaCl or 20 mM imidazole, 1X PBS. Column elutions were concentrated using centrifugal concentrators at 10, 50, or 100 kDa cutoffs, followed by size-exclusion chromatography on an ÄKTA Pure system (Cytiva). ÄKTA pure FPLC with a Superdex 6 Increase (S6) or Superdex 200 Increase (S200) gel filtration column was used for purification. 1 mL of sample was injected using a 2 mL loop and run over the S6 or S200 which had been preequilibrated in degassed 20 mM HEPES, 150 mM NaCl or 1X PBS prior to use and stored at -20°C.

### Fab production and purification

1/10 volume of 1 M Tris, pH 8 was added to IgGs at ∼2 mg/mL in 1X PBS. 2 μL of a 1 mg/mL stock of Lys-C (stock stored at -20°C) was added for each mg of human IgG1 and digested for 1 hour at 37°C with moderate rotation. Digested Fabs were purified using a 5 mL HiTrap SP HP cation exchange chromatography column on an ÄKTA system using 50 mM sodium acetate (NaOAc) pH 5.0 with gradient NaCl elution (using 50 mM NaOAc + 1M NaCl, pH 5.0). Fab fractions were pooled and dialyzed against 1X PBS and concentrated using 30 kDa concentrators. Purified Fabs were stored at -20°C.

### Biolayer interferometry (BLI) binding experiments

All reactions were run on an Octet Red 96 at 30°C and samples were run in 1X PBS with 0.1% BSA and 0.05% Tween 20 (octet buffer). IgGs and Fabs were assessed for binding to biotinylated antigens using streptavidin (SA) biosensors (Sartorius/ForteBio) or to unbiotinylated, His-tagged antigens using Anti-Penta-HIS biosensors (Sartorius/ForteBio). Antigen was loaded to a threshold of 1 nm shift. Tips were then washed and baselined in wells containing only octet buffer. Samples were then associated in wells containing IgG or Fab at 100 nM concentration unless otherwise stated (other concentrations are given in **Supplementary Data 1**). A control well with loaded antigen but that was associated in a well containing only 200 μL octet buffer was used as a baseline subtraction for data analysis. Association and dissociation binding curves were fit in Octet System Data Analysis Software version 9.0.0.15 using a 1:2 bivalent model for IgGs to determine apparent *K*_d_ and a 1:1 model for Fabs to determine *K*_d_. Averages of fitted *K*_d_ values from at least two independent experiments are reported to two significant figures. Wildtype and the highest-affinity variants were also tested at multiple concentrations and *K*_d_ values were averaged across all replicates and concentrations (**Supplementary Data 1**). To estimate measurement error, we compute the coefficient of variation (CV; the ratio of the standard deviation to the mean across replicates) for each antibody-antigen *K*_d_ pair and report the mean CV for each antigen in **Supplementary Tables 2** and **4-9**.

### Thermal melts

We measured thermal melting profiles of proteins by differential scanning fluorimetry on a Prometheus NT.48 instrument. Protein samples (0.1 mg/mL) were loaded into glass capillaries and then subject to a temperature gradient from 20 to 95°C at a heating rate of 1°C per minute. Intrinsic fluorescence (350 nm and 330 nm) was recorded as a function of temperature using PR.ThermControl version 2.3.1 software. Thermal melting curves were plotted using the first derivative of the ratio (350 nm/330 nm). Melting temperatures were calculated automatically by the instrument and represented peaks in the thermal melting curves.

### Lentivirus production

We produced SARS-CoV-2 Spike (D614G and Beta variants) pseudotyped lentiviral particles. Viral transfections were done in HEK293T cells using calcium phosphate transfection reagent. Six million cells were seeded in D10 media [Dulbecco’s Modified Eagle Medium (DMEM) + additives: 10% fetal bovine serum (FBS), L-glutamate, penicillin, streptomycin, and 10 mM HEPES] in 10 cm plates one day prior to transfection. A five-plasmid system was used for viral production, as previously described [58]. The Spike vector contained the 21 amino acid truncated form of the SARS-CoV-2 Spike sequence from the Wuhan-Hu-1 strain of SARS-CoV-2 (GenBank: BCN86353.1) or the Beta variant-of-concern (GenBank: QUT64557.1). The other viral plasmids, used as previously described [58], are pHAGE-Luc2-IRS-ZsGreen (NR-52516), HDM-Hgpm2 (NR-52517), pRC-CMV-Rev1b (NR-52519), and HDM-tat1b (NR-52518). These plasmids were added to D10 medium in the following ratios: 10 μg pHAGE-Luc2-IRS-ZsGreen, 3.4 μg FL Spike, 2.2 μg HDM-Hgpm2, 2.2 μg HDM-Tat1b, 2.2 μg pRC-CMV-Rev1b in a final volume of 1000 μL.

Ebola GP-pseudotyped lentiviruses were produced using the same packaging (pHAGE-Luc2-IRS-ZsGreen) and helper plasmids (HDM-Hgpm2, HDM-Tat1b, pRC-CMV-Rev1b) but with the plasmid encoding full-length Ebola GP (GenBank: AAG40168.1).

After adding plasmids to medium, we added 30 μL BioT (BioLand) to form transfection complexes. Transfection reactions were incubated for 10 minutes at room temperature, and then 9 mL of medium was added slowly. The resultant 10 mL was added to plated HEK cells from which the medium had been removed. Culture medium was removed 24 hours post-transfection and replaced with fresh D10 medium. Viral supernatants were harvested 72 hours post-transfection by spinning at 300X g for five minutes followed by filtering through a 0.45-μm filter. Viral stocks were aliquoted and stored at -80°C until further use.

### Pseudovirus neutralization

The target cells used for infection in SARS-CoV-2 pseudovirus neutralization assays are from a HeLa cell line stably overexpressing human angiotensin-converting enzyme 2 (ACE2), as well as the protease known to process SARS-CoV-2, transmembrane serine protease 2 (TMPRSS2). Production of this cell line is described in detail previously [59], with the addition of stable TMPRSS2 incorporation. ACE2/TMPRSS2/HeLa cells were plated one day prior to infection at 8,000 cells per well. For Ebola-pseudovirus neutralization assays, HEK-293T cells were seeded in 96-well plates one day prior to infection at 20,000 cells per well. 96-well, white-walled, white-bottom plates were used for neutralization assays (Thermo Fisher Scientific).

On the day of the assay, purified IgGs in 1X PBS were sterile filtered using a 0.22-μm filter. Dilutions of this filtered stock were made into sterile 1X Dulbecco’s PBS (DPBS) (Thermo Fisher Scientific) which was 5% by volume D10 medium. Samples were run in technical quadruplicate in each experiment. A virus mixture was made containing the virus of interest (for example SARS-CoV-2) and D10 media (DMEM + additives: 10% FBS, L-glutamate, penicillin, streptomycin, and 10 mM HEPES). Virus dilutions into media were selected such that a suitable signal would be obtained in the virus-only wells. A suitable signal was selected such that the virus only wells would achieve a luminescence of at least >5,000,000 relative light units (RLU). 60 μL of this virus mixture was added to each of the antibody dilutions to make a final volume of 120 μL in each well. Virus-only wells were made which contained 60 μL 1X DPBS and 60 μL virus mixture. Cells-only wells were made which contained 120 μL D10 media.

The antibody/virus mixture was left to incubate for 1 hour at 37°C. Following incubation, the medium was removed from the cells on the plates made 1 day prior. This was replaced with 100 μL of antibody/virus dilutions and incubated at 37°C for approximately 24 hours. Infectivity readout was performed by measuring luciferase levels. SARS-CoV-2 and Ebola pseudovirus neutralization assays were read out 48- and 72-hour post-infection, respectively. Medium was removed from all wells and cells were lysed by the addition of 100 μL BriteLite™ assay readout solution (Perkin Elmer) into each well. Luminescence values were measured using an Infinite^®^ 200 PRO Microplate Reader (Tecan) using i-control version 2.0 software (Tecan). Each plate was normalized by averaging the cells-only (0% infection) and virus-only (100% infection) wells. We used the neutcurve Python package version 0.5.7 to fit the normalized datapoints and to compute the IC_50_ values, which we report to two significant digits. To estimate measurement error, we compute the CV for each antibody-virus IC_50_ pair and report the mean CV for each virus in **Supplementary Tables 5, 8,** and **9**.

### Computing frequency of changes to antibody protein sequences

We computed the frequency of residues involved in affinity-enhancing substitutions by aligning the wildtype VH and VL sequences of our antibodies to databases of protein sequences. The first database we considered is UniRef90, where we use the same database release used to train ESM-1v. For each antibody protein sequence, we obtained the set of 2,000 sequences in the database that are closest to the antibody by sequence similarity based on Levenshtein distance (with the farthest sequences having between 24% to 49% sequence similarity); we compute sequence similarity using the fuzzywuzzy Python package version 0.18.0. We then use mafft version 7.475 to perform multiple sequence alignment among the set of sequences. We used the alignment to compute amino acid frequencies at each site in the VH or VL sequence. The second database we considered is provided by the abYsis webtool. We aligned VH and VL protein sequences using the default settings provided in the “Annotate” tool, using the database of “All” sequences as of March 1, 2022. While our frequency estimation procedure is based on the entire UniRef90 dataset, we also sought to count the number of annotated immunoglobulin variable regions in UniRef90; to do so, we used the UniRef query tool (https://www.uniprot.org/uniref/) with the queries “name:“immunoglobulin heavy variable” AND identity:0.9”, “name:“immunoglobulin kappa variable” AND identity:0.9”, and “name:“immunoglobulin lambda variable” AND identity:0.9”.

### Natural protein evaluation based on scanning mutagenesis data

We evaluated the ability for the language models and algorithms used in our study to guide efficient evolution in other settings beyond antibodies. We leveraged deep mutational scanning (DMS) datasets to validate that our approach would enable a researcher to acquire high-fitness variants. We used all DMS datasets from the benchmarking study by Livesey and Marsh [41] with 90% or higher coverage of all single-residue substitutions; variants that were not measured were treated as having low fitness. We also used a scanning mutagenesis dataset generated by Markin et al. that measured Michaelis-Menten kinetics of all single-site glycine or valine substitutions to the bacterial enzyme PafA; for this dataset, any language-model-recommended substitutions that did not involve glycine or valine substitutions were excluded from the analysis. We applied a cutoff for each dataset to binarize sequences as high- or low-fitness variants (cutoffs are provided in **Supplementary Table 11**); we then compared enrichment of high-fitness variants among the language-model-recommended variants to the background frequency of high-fitness variants among all single-residue substitutions. For these proteins, as with our antibody experiments, we chose values of *k* that result in a small number (∼10^1^) of acquired substitutions: we used *α* = 1 and *k* = 2 for all proteins except those where this resulted in |𝒜| < 5, in which case we set *k* = 1.

To quantify the statistical significance of an enrichment, we assumed that the null distribution of the number of high-fitness, language-model-recommended variants was given by a hypergeometric distribution parameterized by the number of language-model-recommended variants |𝒜|, the number of high-fitness variants among the all single-residue substitutions, and the total number of single-residue substitutions considered, which we used to compute a one-sided *P* value. We used the hypergeometric calculator at https://stattrek.com/online-calculator/hypergeometric.aspx.

To test the relationship between likelihood stringency and the fraction of high-fitness substitutions, we also performed a small-scale parameter sweep varying the cutoff values *α* and *k* and computing: (1) the percentage fraction of high-fitness substitutions in 𝒜, (2) the maximum fitness value of a variant in 𝒜 divided by the maximum fitness value of a variant across the full mutational scan, and (3) the maximum fitness value of a variant in 𝒜 divided by the 99^th^ percentile of the fitness values across the full mutational scan; prior to this normalization, the raw fitness values are also linearly scaled to take values between 0 and 1, inclusive. Normalized values, the number of acquired variants |𝒜|, and the parameter combinations are plotted in **Supplementary Fig. 2**.

### Data availability

Raw data for this study has been deposited to Zenodo at DOI:10.5281/zenodo.6415457. *K*_d_, IC_50_, and *T*_m_ values across replicate experiments are available as **Supplementary Data 1**.

### Code availability

Code and scripts used in this study are available at https://github.com/brianhie/efficient-evolution.

## Acknowledgements

We thank Benjamin Bell and Ashwin Narayan for helpful discussions. We thank the Peter Kim Lab for useful comments on the manuscript. B.L.H. acknowledges the support of the Stanford Science Fellows program. This work was supported by the Virginia & D.K. Ludwig Fund for Cancer Research (P.S.K.) and by the Chan Zuckerberg Biohub (P.S.K.).

## Author contributions

Conceptualization, investigation, and interpretation: B.L.H. and P.S.K. Computational experiments and software development: B.L.H. Antibody cloning, expression, and purification: B.L.H. and V.R.S. Antigen cloning, expression, and purification: B.L.H., D.X., T.U.J.B., P.A.W., and S.T. Fab production: B.L.H. Binding assays: B.L.H. Thermal melts: B.L.H. and V.R.S. Lentivirus production: D.X. Pseudovirus neutralization: D.X. Writing (initial draft): B.L.H. Writing (final draft): all authors.

**Supplementary Data** (enclosure)

**Supplementary Data 1:** Experimental *K*_d_, IC_50_, and *T*_m_ values across seven antibody directed evolution campaigns.

## Supplementary Tables, Figures, and Information

**Supplementary Table 1:** Summary of antibodies considered in this study. Information on the antibodies considered in each of our directed evolution campaigns. Matured indicates extensive somatic hypermutation from germline (and, in the case of MEDI8852, additional in-vitro affinity maturation). Source indicates how the antibody sequence was obtained; germline-inferred sequences were obtained from the original publications. Improved binding is defined as a 1.1-fold improvement or higher from wildtype. Preserved binding is defined as a sub-micromolar *K*_d_ for the screened antigen. ND: not determined.

| Antibody | Matured? | Source | Virus | Screened antigen | Round-1 variants with improved binding | Round-1 variants with preserved binding | Best affinity improvement (antigen) | Best neutralization improvement (virus) |
| --- | --- | --- | --- | --- | --- | --- | --- | --- |
| MEDI8852 [27] | Yes (in vivo + in vitro) | Patient-derived + in vitro evolution | Influenza A | HA H4 Hubei | 2 / 13 = 15% | 13 / 14 = 93% | 7-fold (HA H7 HK17) | ND |
| MEDI8852 UCA [27] | No | Germline-inferred | Influenza A | HA H1 Solomon | 3 / 8 = 38% | 8 / 8 = 100% | 23-fold (HA H4 Hubei) | ND |
| mAb114 [28] | Yes | Patient-derived | Ebolavirus | GP | 7 / 13 = 54% | 13 / 13 = 100% | 3.4-fold (GP) | 1.5-fold (GP pseudotyped lentivirus) |
| mAb114 UCA [27] | No | Germline-inferred | Ebolavirus | GP | 6 / 9 = 67% | 7 / 9 = 78% | 160-fold (GP) | >100-fold (GP pseudotyped lentivirus) |
| S309 [29] | Yes | Patient-derived | Sarbecovirus | Wuhan-Hu-1 S-6P | 2 / 10 = 20% | 9 / 10 = 90% | 1.7-fold (Beta S-6P) | ND |
| REGN10987 [30] | Yes | Patient-derived | SARS-CoV-2 | Beta S-6P | 2 / 8 = 25% | 8 / 8 = 100% | 1.3-fold (Beta S-6P) | 2-fold (Beta pseudotyped lentivirus) |
| C143 [36] | No | Patient-derived | SARS-CoV-2 | Beta S-6P | 10 / 14 = 71% | 10 / 14 = 71% | 13-fold (Beta S-6P) | 31-fold (Beta pseudotyped lentivirus) |

**Supplementary Table 2:** MEDI8852 variants. Variants tested across two rounds of directed evolution, the corresponding Kabat-annotated regions, and *K*_d_ values for three HA antigens. The wildtype row is highlighted in gray; variants with improved affinity are highlighted in blue. NB: no binding; ND: not determined; CV: coefficient of variation.

| Round | Design | Region | H4 Hubei<br>Fab $K_d$ (nM) | H7 HK16<br>Fab $K_d$ (nM) | H7 HK17<br>Fab $K_d$ (nM) |
| --- | --- | --- | --- | --- | --- |
|  | MEDI8825<br>WT |  | 0.56 | 6.2 | 0.21 |
|  | VH D27F | HFR1 | 0.90 | 9.0 | ND |
|  | VH V35Y | CDR-H1 | 0.87 | 8.6 | ND |
|  | VH S44G | HFR2 | 0.60 | 7.0 | ND |
|  | VH T53I | CDR-H2 | 0.75 | 10 | ND |
|  | VH W59S | CDR-H2 | 0.78 | 7.1 | ND |
|  | VH E65P | CDR-H2 | 0.36 | 2.7 | 0.10 |
|  | VH N74S | HFR3 | 0.53 | 6.0 | ND |
|  | VH M117Y | CDR-H3 | 0.36 | 4.1 | ND |
|  | VL T25A | CDR-L1 | 1.4 | 17 | ND |
|  | VL L29V | CDR-L1 | 4.2 | 80 | ND |
|  | VL T33L | CDR-L1 | 3.2 | 41 | ND |
|  | VL G55A | CDR-L2 | 0.51 | 5.1 | ND |
|  | VL R92D | CDR-L3 | NB | 26 | ND |
|  | VL G95P | LFR4 | 0.80 | 5.3 | ND |
| Round 2 | VH E65P-M117Y | Multi | 0.31 | 2.5 | 0.030 |
| Error (mean CV): | | | $\pm 12\%$ | $\pm 14\%$ | $\pm 34\%$ |

**Supplementary Table 3:** Binding to Group 1 and Group 2 HAs. Binding affinity between MEDI8852 WT Fab and three variant Fabs against a panel of nine HAs. A *K*_d_ of <0.001 indicates an interaction with no observed dissociation when measured via BLI.

| Antigen<br>(Group) | Antibody | Fab $K_d$ (nM) |
| --- | --- | --- |
| H1 Caledonia<br>(Group 1) | MEDI8852 | <0.001 |
|  | VH E65P | <0.001 |
|  | VH E65P-M117Y | <0.001 |
| H1 Solomon<br>(Group 1) | MEDI8852 | <0.001 |
|  | VH E65P | <0.001 |
|  | VH E65P-M117Y | <0.001 |
| H2 Japan<br>(Group 1) | MEDI8852 | <0.001 |
|  | VH E65P | <0.001 |
|  | VH E65P-M117Y | <0.001 |
| H3 Panama<br>(Group 2) | MEDI8852 | <0.001 |
|  | VH E65P | <0.001 |
|  | VH E65P-M117Y | <0.001 |
| H3 Victoria<br>(Group 2) | MEDI8852 | <0.001 |
|  | VH E65P | <0.001 |
|  | VH E65P-M117Y | <0.001 |
| H4 Hubei<br>(Group 2) | MEDI8852 | 0.56 |
|  | VH E65P | 0.36 |
|  | VH E65P-M117Y | 0.31 |
| H5 Vietnam<br>(Group 1) | MEDI8852 | <0.001 |
|  | VH E65P | <0.001 |
|  | VH E65P-M117Y | <0.001 |
| H7 HK16<br>(Group 2) | MEDI8852 | 6.2 |
|  | VH E65P | 2.7 |
|  | VH E65P-M117Y | 2.5 |
| H7 HK17<br>(Group 2) | MEDI8852 | 0.21 |
|  | VH E65P | 0.10 |
|  | VH E65P-M117Y | 0.030 |

**Supplementary Table 4:** MEDI8852 UCA variants. Variants tested across two rounds of directed evolution, the corresponding Kabat-annotated regions, and *K*_d_ values for three HA antigens. In Round 2, all possible combinations involving K58S, V65P, P75R in the VH and G95P in the VL were made. The wildtype row is highlighted in gray; variants with improved affinity are highlighted in blue. *K*_d_ app.: *K*_d_ apparent; ND: not determined; CV: coefficient of variation.

| Round | Design | Region | H1<br>Solomon<br>IgG $K_d$<br>app. (nM) | H1<br>Solomon<br>Fab $K_d$<br>(nM) | H4 Hubei<br>IgG $K_d$<br>app. (nM) | H4 Hubei<br>Fab $K_d$<br>(μM) | H7 HK17<br>$K_d$ (nM) |
| --- | --- | --- | --- | --- | --- | --- | --- |
| | MEDI8852 UCA WT | | 2.0 | 140 | 740 | 29 | IgG app. $K_d$:<br>17<br>Fab $K_d$:<br>4400 |
| Round 1 | VH S44G | HFR2 | 2.1 | ND | ND | ND | ND |
|  | VH T53I | CDR-H2 | 2.1 | ND | ND | ND | ND |
|  | VH K58S | CDR-H2 | 1.5 | 110 | ND | 2.5 | ND |
|  | VH V65P | CDR-H2 | 1.7 | ND | ND | ND | ND |
|  | VH N74S | HFR3 | 2.6 | ND | ND | ND | ND |
|  | VH P75R | HFR3 | 2.0 | ND | ND | ND | ND |
|  | VL N34A | CDR-L1 | 8.2 | ND | ND | ND | ND |
|  | VL G95P | LFR4 | 1.3 | 65 | ND | 17 | ND |
| Round 2 | VH K58S-V65P | CDR-H2 | 1.3 | ND | 80 | ND | ND |
|  | VH K58S-P75R | Multi | 1.2 | ND | 47 | ND | ND |
|  | VH V65P-P75R | Multi | 1.5 | ND | 570 | ND | ND |
|  | VH K58S-V65P-P75R | Multi | 1.2 | ND | 72 | ND | ND |
|  | VH K58S / VL G95P | Multi | 0.96 | 56 | 36 | 3.1 | ND |
|  | VH V65P / VL G95P | Multi | 0.93 | ND | 100 | ND | ND |
|  | VH P75R / VL G95P | Multi | 0.95 | ND | 110 | ND | ND |
| | VH K58S-V65P / VL G95P | Multi | 0.78 | 50 | 30 | 1.3 | IgG app. $K_d$:<br>5.6<br>Fab $K_d$:<br>810 |
|  | VH K58S-P75R / VL G95P | Multi | 0.88 | ND | 45 | ND | ND |
|  | VH V65P-P75R / VL G95P | Multi | 0.87 | ND | 110 | ND | ND |
|  | VH K58S-V65P-P75R / VL G95P | Multi | 0.79 | 55 | 42 | 1.5 | ND |
| Error (mean CV): | | | $\pm 11\%$ | $\pm 8.6\%$ | $\pm 10\%$ | $\pm 16\%$ | $\pm 13\%$ |

**Supplementary Table 5:** mAb114 variants. Variants tested across two rounds of directed evolution, the corresponding Kabat-annotated regions, and *K*_d_ values for ebolavirus GP. Neutralization IC_50_ values were also determined for mAb114 WT and all Round-2 variant IgGs against GP-pseudotyped lentivirus. The wildtype row is highlighted in gray; variants with improved affinity are highlighted in blue. ND: not determined; CV: coefficient of variation.

| Round | Design | Region | GP Fab $K_d$ (nM) | Ebola GP pseudotyped lentivirus $IC_{50}$ (nM) |
| --- | --- | --- | --- | --- |
|  | mAb114 WT |  | 0.21 | 1.4 |
| Round 1 | VH M31S | CDR-H1 | 0.13 | ND |
|  | VH I41P | HFR2 | 0.33 | ND |
|  | VH D42G | HFR2 | 0.18 | ND |
|  | VH A68T | HFR3 | 0.10 | ND |
|  | VH E72D | HFR3 | 0.13 | ND |
|  | VH S79Y | HFR3 | 0.12 | ND |
|  | VH I113T | HFR4 | 0.071 | ND |
|  | VL I19V | LFR1 | 0.23 | ND |
|  | VL F29I | CDR-L1 | 0.46 | ND |
|  | VL V43A | LFR2 | 0.068 | ND |
|  | VL S49Y | LFR2 | 0.58 | ND |
|  | VL H70D | LFR3 | 0.27 | ND |
|  | VL N90Q | CDR-L3 | 4.0 | ND |
| Round 2 | VH D42G-A68T-S79Y | Multi | 0.19 | 1.2 |
|  | VH I41P-D42G-A68T-S79Y-I113T | Multi | 0.19 | 1.2 |
|  | VH A68T-I113T / VL V43A | Multi | 0.17 | 1.1 |
|  | VH A68T-E72D-S79Y-I113T / VL V43A | Multi | 0.076 | 1.2 |
|  | VH D42G-A68T-S79Y / VL V43A | Multi | 0.061 | 0.96 |
|  | VH I41P-D42G-A68T-S79Y-I113T / VL V43A | Multi | 0.069 | 1.0 |
| Error (mean CV): | | | $\pm 41\%$ | $\pm 18\%$ |

**Supplementary Table 6:**
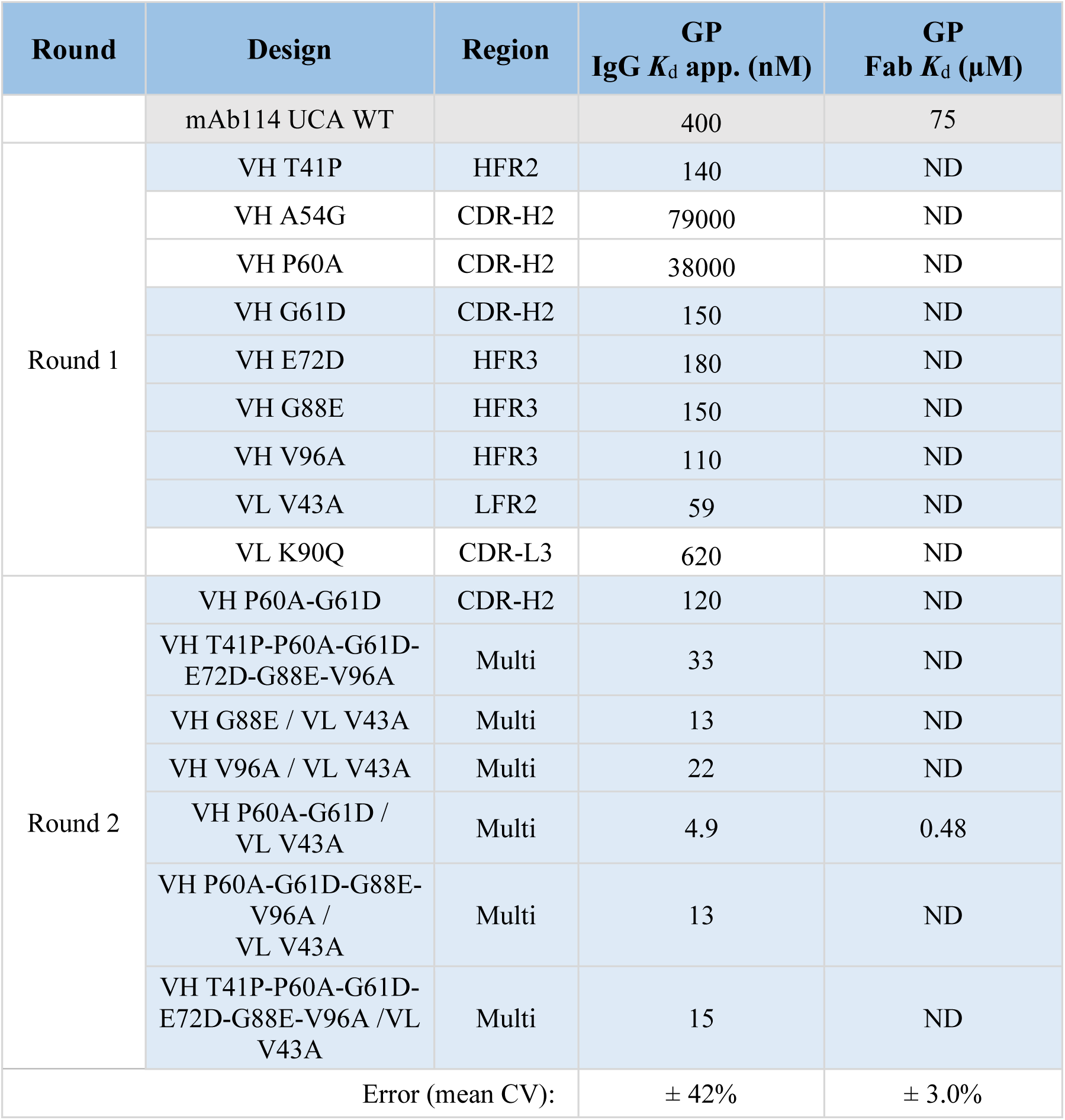
mAb114 UCA variants. Variants tested across two rounds of directed evolution, the corresponding Kabat-annotated regions, and *K*_d_ values (for both IgG and Fab versions) for ebolavirus GP. The wildtype row is highlighted in gray; variants with improved affinity are highlighted in blue. *K*_d_ app.: *K*_d_ apparent; ND: not determined; CV: coefficient of variation.

| Round | Design | Region | GP<br>IgG $K_d$ app. (nM) | GP<br>Fab $K_d$ (μM) |
| --- | --- | --- | --- | --- |
|  | mAb114 UCA WT |  | 400 | 75 |
| Round 1 | VH T41P | HFR2 | 140 | ND |
|  | VH A54G | CDR-H2 | 79000 | ND |
|  | VH P60A | CDR-H2 | 38000 | ND |
|  | VH G61D | CDR-H2 | 150 | ND |
|  | VH E72D | HFR3 | 180 | ND |
|  | VH G88E | HFR3 | 150 | ND |
|  | VH V96A | HFR3 | 110 | ND |
|  | VL V43A | LFR2 | 59 | ND |
|  | VL K90Q | CDR-L3 | 620 | ND |
| Round 2 | VH P60A-G61D | CDR-H2 | 120 | ND |
|  | VH T41P-P60A-G61D-E72D-G88E-V96A | Multi | 33 | ND |
|  | VH G88E / VL V43A | Multi | 13 | ND |
|  | VH V96A / VL V43A | Multi | 22 | ND |
|  | VH P60A-G61D /<br>VL V43A | Multi | 4.9 | 0.48 |
|  | VH P60A-G61D-G88E-V96A /<br>VL V43A | Multi | 13 | ND |
|  | VH T41P-P60A-G61D-E72D-G88E-V96A /VL<br>V43A | Multi | 15 | ND |
| Error (mean CV): | | | $\pm 42\%$ | $\pm 3.0\%$ |

**Supplementary Table 7:** S309 variants. Variants tested across two rounds of directed evolution, the corresponding Kabat-annotated regions, and *K*_d_ values for antigens from three SARS-CoV-2 variants. The wildtype row is highlighted in gray; variants with improved affinity are highlighted in blue. W1: Wuhan-Hu-1; NE: no expression; ND: not determined; CV: coefficient of variation.

| Round | Design | Region | W1<br>S-6P Fab<br>$K_d$ (nM) | Beta S-6P<br>Fab $K_d$<br>(nM) | Omicron<br>RBD Fab<br>$K_d$ (nM) |
| --- | --- | --- | --- | --- | --- |
|  | S309 WT |  | 2.5 | 2.4 | 14 |
|  | Sotrovimab<br>(VH N55Q) | CDR-H2 | 2.4 | 2.0 | 17 |
| Round 1 | VH P28T | HFR1 | 4.3 | ND | ND |
|  | VH T77N | HFR3 | 2.8 | ND | ND |
|  | VH G79A | HFR3 | 2.8 | ND | ND |
|  | VH R84S | HFR3 | 2.7 | ND | ND |
|  | VH R85S | HFR3 | 3.1 | ND | ND |
|  | VH R87T | HFR3 | 2.1 | ND | ND |
|  | VL T28S | CDR-L1 | 2.1 | ND | ND |
|  | VL T32N | CDR-L1 | 2.7 | ND | ND |
|  | VL S95V | CDR-L3 | NE | ND | ND |
|  | VL L96P | CDR-L3 | 2.7 | ND | ND |
| Round 2 | VH T77N-G79A-R84S | HFR3 | 3.2 | ND | ND |
|  | VH T77N-G79A-R84S-R85S | HFR3 | 3.1 | ND | ND |
|  | VL T28S-T32N | CDR-L1 | 3.0 | ND | ND |
|  | VH G79A /<br>VL T28S | Multi | 2.0 | 1.4 | 15 |
|  | VH R84S / VL T28S | Multi | 2.5 | ND | ND |
|  | VH R87T / VL T28S | Multi | 2.5 | ND | ND |
|  | VH T77N-G79A-R84S<br>/ VL T28S | Multi | 3.3 | ND | ND |
|  | VH T77N-G79A-R84S<br>/ VL T28S-T32N | Multi | 3.5 | ND | ND |
|  | VH T77N-G79A-R84S-R85S /<br>VL T28S-T32N | Multi | 4.3 | ND | ND |
| Error (mean CV): |  |  | ± 19% | ± 14% | ± 16% |

**Supplementary Table 8:** REGN10987 variants. Variants tested across two rounds of directed evolution, the corresponding Kabat-annotated regions, and *K*_d_ values for antigens from two SARS-CoV-2 variants. Round-1 variants were pre-screened as IgGs with a single replicate before testing the highest-avidity variants as Fabs and with multiple replicates. Neutralization IC_50_ values were also determined for REGN10987 WT and selected affinity-enhancing variant IgGs against D614G- and Beta-pseudotyped lentivirus. The wildtype row is highlighted in gray; variants with improved affinity are highlighted in blue. *K*_d_ app.: *K*_d_ apparent; NA: not applicable; ND: not determined; CV: coefficient of variation.

| Round | Design | Region | Beta S-6P<br>IgG $K_d$<br>app. (nM) | Beta S-6P<br>Fab $K_d$<br>(nM) | Omicron<br>RBD IgG<br>$K_d$ app.<br>(μM) | D614G-pseudotyped<br>lentivirus IC<sub>50</sub><br>(ng/mL) | Beta-pseudotyped<br>lentivirus IC<sub>50</sub><br>(ng/mL) |
| --- | --- | --- | --- | --- | --- | --- | --- |
| | REGN10987 WT | | 0.091 | 13 | 0.70<br>(Fab $K_d$:<br>23) | 1.1 | 1.4 |
| Round 1 | VH R16G | HFR1 | 0.073 | 10 | 0.51 | 0.83 | 0.73 |
|  | VH S98R | HFR3 | 0.41 | ND | ND | ND | ND |
|  | VH V108D | CDR-H3 | 0.076 | 26 | ND | ND | ND |
|  | VL S82A | LFR3 | 0.063 | 14 | ND | ND | ND |
|  | VL N91C | CDR-L3 | 0.053 | 12 | ND | ND | ND |
|  | VL N91S | CDR-L3 | 0.066 | 22 | ND | ND | ND |
|  | VL L93Y | CDR-L3 | 0.30 | ND | ND | ND | ND |
|  | VL I96S | CDR-L3 | 0.072 | 15 | ND | ND | ND |
| Round 2 | VH R16G-V108D | Multi | ND | 30 | 2.2 | ND | ND |
|  | VL S82A-N91C-I96S | Multi | ND | 12 | 0.48 | 1.0 | 0.87 |
|  | VH R16G /<br>VL S82A | Multi | ND | 12 | 0.88 | 1.0 | 0.78 |
| | VH R16G /<br>VL N91C | Multi | ND | 30 | 0.42<br>(Fab $K_d$:<br>4.5) | ND | ND |
|  | VH R16G / VL I96S | Multi | ND | 12 | 1.2 | 1.2 | 0.79 |
|  | VH R16G-V108D /<br>VL S82A-N91C-I96S | Multi | ND | 40 | 2.5 | ND | ND |
| Error (mean CV): | | | NA | ± 30% | ± 19% | ± 30% | ± 15% |

**Supplementary Table 9:**
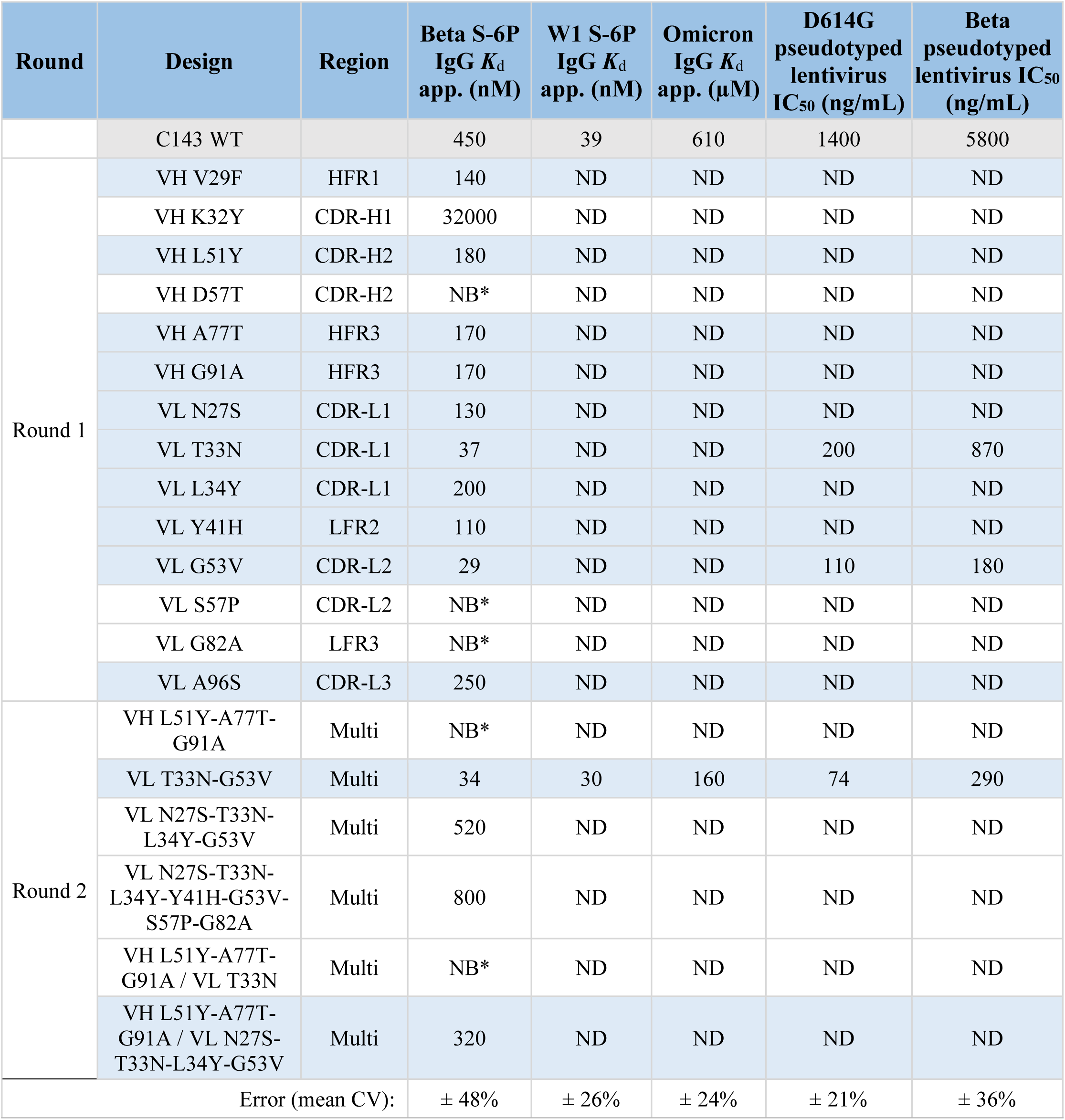
C143 variants. Variants tested across two rounds of directed evolution, the corresponding Kabat-annotated regions, and *K*_d_ values for antigens from three SARS-CoV-2 variants. Neutralization IC_50_ values were also determined for C143 WT and selected affinity-enhancing variant IgGs against D614G- and Beta-pseudotyped lentivirus. The wildtype row is highlighted in gray; variants with improved affinity are highlighted in blue. An asterisk (*) indicates examples where binding was observed but BLI data were not suitable for fitting. W1: Wuhan-Hu-1; *K*_d_ app.: *K*_d_ apparent; NB: no binding; ND: not determined; CV: coefficient of variation.

**Supplementary Table 10:** Originality of affinity-enhancing substitutions. Each row corresponds to an amino acid substitution that enhances the binding affinity of its corresponding variant antibody, and some of which also enhance affinity in combination with other substitutions. We computed frequencies of amino acid substitutions among natural sequences using two datasets, UniRef90 and abYsis (**Methods**); UniRef90 was the sequence database used to train the language models in our algorithm and abYsis is a separate, curated database of natural antibody sequences. The “wildtype residue frequency” indicates the percentage of sequences in a multiple sequence alignment with the same residue as wildtype at the given position; the “mutant residue frequency” is the same statistic except for the mutant residue. The “top residue” indicates the amino acid with the highest frequency observed at the given site, the “top residue frequency” indicates the percentage of sequences that contain the top residue at the given site, and dashes indicate settings in which the mutant residue is also the top residue. Substitutions with frequencies up to 5% are considered “rare,” those with frequencies above 5% and up to 10% are considered “uncommon,” and those above 10% are considered “common.” Blue shading indicates substitutions to rare or uncommon residues according to frequency information from either UniRef90 or abYsis.

| Antibody | Affinity-enhancing substitution | UniRef90 (training dataset) |  |  |  |  | abYsis |  |  |  |
| --- | --- | --- | --- | --- | --- | --- | --- | --- | --- | --- |
|  |  | Wildtype residue frequency | Mutant residue frequency | Top residue | Top residue frequency | Notes | Wildtype residue frequency | Mutant residue frequency | Top residue | Top residue frequency |
| MEDI8852 | VH E65P | 6% | 32% | D | 35% |  | 14% | 22% | D | 32% |
|  | VH M117Y | <1% | 59% | - | - |  | <1% | 49% | - | - |
| MEDI8852 UCA | VH K58S | 8% | 3% | T | 67% | Uncommon to rare | 2% | 30% | G | 35% |
|  | VH V65P | 1% | 32% | D | 36% |  | 2% | 22% | D | 32% |
|  | VL G95P | 44% | 2% | G | 44% | Common to rare | 99% | <1% | G | 99% |
| mAb114 | VH M31S | <1% | 45% | - | - |  | <1% | 48% | - | - |
|  | VH D42G | 1% | 88% | - | - |  | <1% | 91% | - | - |
|  | VH A68T | 1% | 88% | - | - |  | 1% | 85% | - | - |
|  | VH E72D | 4% | 91% | - | - |  | 2% | 93% | - | - |
|  | VH S79Y | 8% | 74% | - | - |  | 16% | 67% | - | - |
|  | VH I113T | 2% | 69% | - | - |  | 2% | 89% | - | - |
|  | VL V43A | 4% | 50% | - | - |  | 2% | 57% | - | - |
| mAb114 UCA | VH T41P | 1% | 94% | - | - |  | 1% | 92% | - | - |
|  | VH G61D | 2% | 57% | - | - |  | 1% | 32% | - | - |
|  | VH E72D | 4% | 92% | - | - |  | 2% | 93% | - | - |
|  | VH G88E | 1% | 85% | - | - |  | 2% | 67% | - | - |
|  | VH V96A | 6% | 77% | - | - |  | 5% | 85% | - | - |
|  | VL V43A | 4% | 50% | - | - |  | 2% | 57% | - | - |
| S309 | VH R87T | 37% | 30% | R | 37% |  | 39% | 41% | - | - |
|  | VL T28S | 1% | 54% | - | - |  | 2% | 77% | - | - |
| REGN10987 | VH R16G | 5% | 68% | - | - |  | 6% | 31% | - | - |
|  | VL N91C | 2% | 2% | Q | 39% | Rare to rare | 1% | 4% | Q | 52% |
| C143 | VH V29F | 3% | 78% | - | - |  | 4% | 69% | - | - |
|  | VH L51Y | 4% | 18% | - | - |  | 1% | <1% | I | 85% |
|  | VH A77T | 1% | 56% | - | - |  | <1% | 62% | - | - |
|  | VH G91A | 4% | 95% | - | - |  | 2% | 97% | - | - |
|  | VL N27S | 11% | 53% | - | - |  | 3% | 77% | - | - |
|  | VL T33N | 2% | 44% | - | - |  | 2% | 36% | - | - |
|  | VL L34Y | 4% | 34% | - | - |  | 2% | 59% | - | - |
|  | VL Y41H | <1% | 6% | K | 59% | Rare to uncommon | <1% | 5% | K | 67% |
|  | VL G53V | 2% | 10% | A | 30% | Rare to uncommon | 4% | 12% | A | 40% |
|  | VL A96S | 2% | 57% | - | - |  | 2% | 36% | - | - |

**Supplementary Table 11:** Enrichment of high-fitness variants based on language-model-recommended substitutions. Each row corresponds to a protein tested via a high-throughput scanning mutagenesis assay that measures various notions of protein fitness, which are summarized in the “Fitness setting” column. All assays involve deep mutational scans that profile variants that represent 90% or more coverage of all single-residue substitutions except for that of PafA, which changes every residue to either a glycine or a valine. The cutoff indicates the study-specific criterion for determining a high-fitness variant. The “Sample size” indicates the number of acquired variants (|𝒜|) and “Sample successes” indicates the number of those variants with high fitness according to the cutoff. The “Population size” indicates the number of variants profiled in the scanning mutagenesis assay, where “Population successes” indicates the number of those variants with high fitness according to the cutoff. “Hit rate” indicates the percentage fraction of high-fitness variants among the language-model-recommended variants (sample successes divided by sample size) whereas “Background” indicates the percentage fraction of high-fitness variants among all single-residue variants (population successes divided by population size). The hypergeometric *P* value computes enrichment of high-fitness variants among the acquired variants by assuming that the number of sample successes has a hypergeometric null distribution with parameters given by the other values (sample size, population successes, and population size); blue shading indicates a one-sided, hypergeometric *P*-value of less than 0.05.

| Protein | Reference | Organism | Fitness setting | Experiment | Cutoff | Sample successes | Sample size | Population successes | Population size | Hit rate (%) | Background (%) | Hypergeometric *P* |
| --- | --- | --- | --- | --- | --- | --- | --- | --- | --- | --- | --- | --- |
| ADRB2 | [60] | Human | Signal transduction + pathway reporter | Deep mutational scan | > 2.8 | 2 | 9 | 915 | 7847 | 22.2 | 11.7 | 0.28 |
| β-lactamase | [61] | Bacteria | Antibiotic resistance (ampicillin, 2500 ug/mL) | Deep mutational scan | > 0.01 | 4 | 10 | 394 | 5434 | 40.0 | 7.2 | 0.0040 |
| Env | [62] | Virus | Viral replication fitness | Deep mutational scan | > 0.1 | 7 | 37 | 749 | 16359 | 18.9 | 4.6 | 0.0013 |
| HA H1 | [63] | Virus | Viral replication fitness | Deep mutational scan | > 0.1 | 5 | 32 | 646 | 10736 | 15.6 | 6.0 | 0.041 |
| HA H3 | [64] | Virus | Viral replication fitness | Deep mutational scan | > 0.1 | 5 | 16 | 715 | 10754 | 31.3 | 6.6 | 0.0030 |
| infA | [65] | Bacteria | Competitive growth | Deep mutational scan | > 0.98 | 4 | 10 | 306 | 1369 | 40.0 | 22.4 | 0.17 |
| MAPK1 | [66] | Human | Competitive growth (SCH772984) | Deep mutational scan | > 2.5 | 1 | 13 | 77 | 6810 | 7.7 | 1.1 | 0.14 |
| P53 | [67] | Human | Competitive growth (etoposide) | Deep mutational scan | > 1 | 2 | 17 | 906 | 7467 | 11.8 | 12.1 | 0.63 |
| PafA | [42] | Bacteria | $K_{cat}/K_M$ | Scanning mutagenesis | $P < 0.01$, faster than WT | 2 | 10 | 35 | 1041 | 20.0 | 3.4 | 0.042 |

**Supplementary Fig. 1:**
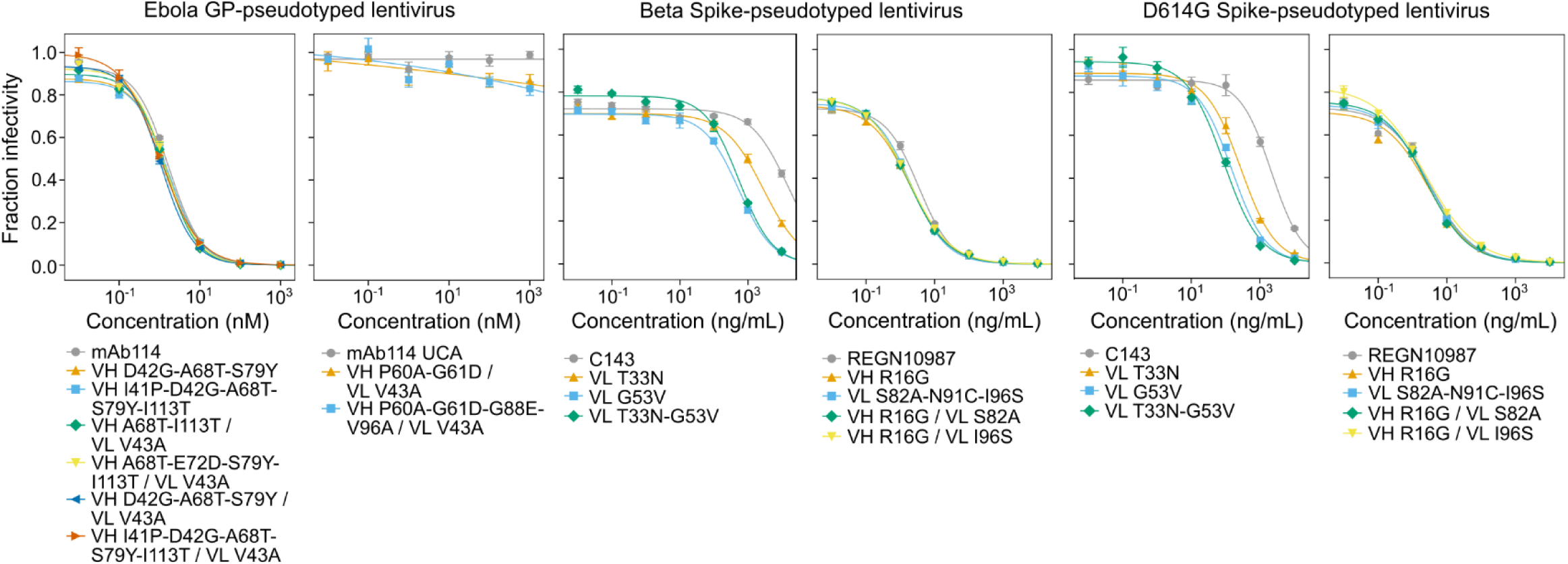
Pseudovirus neutralization of affinity-matured variants. Neutralization curves for wildtype antibodies (gray) and variants obtained by our language-model-guided affinity maturation campaigns. Also see **Supplementary Tables 5, 8**, and **9** for corresponding IC_50_ values. Points indicate the mean; error bars indicate the standard deviation.

**Supplementary Fig. 2:**
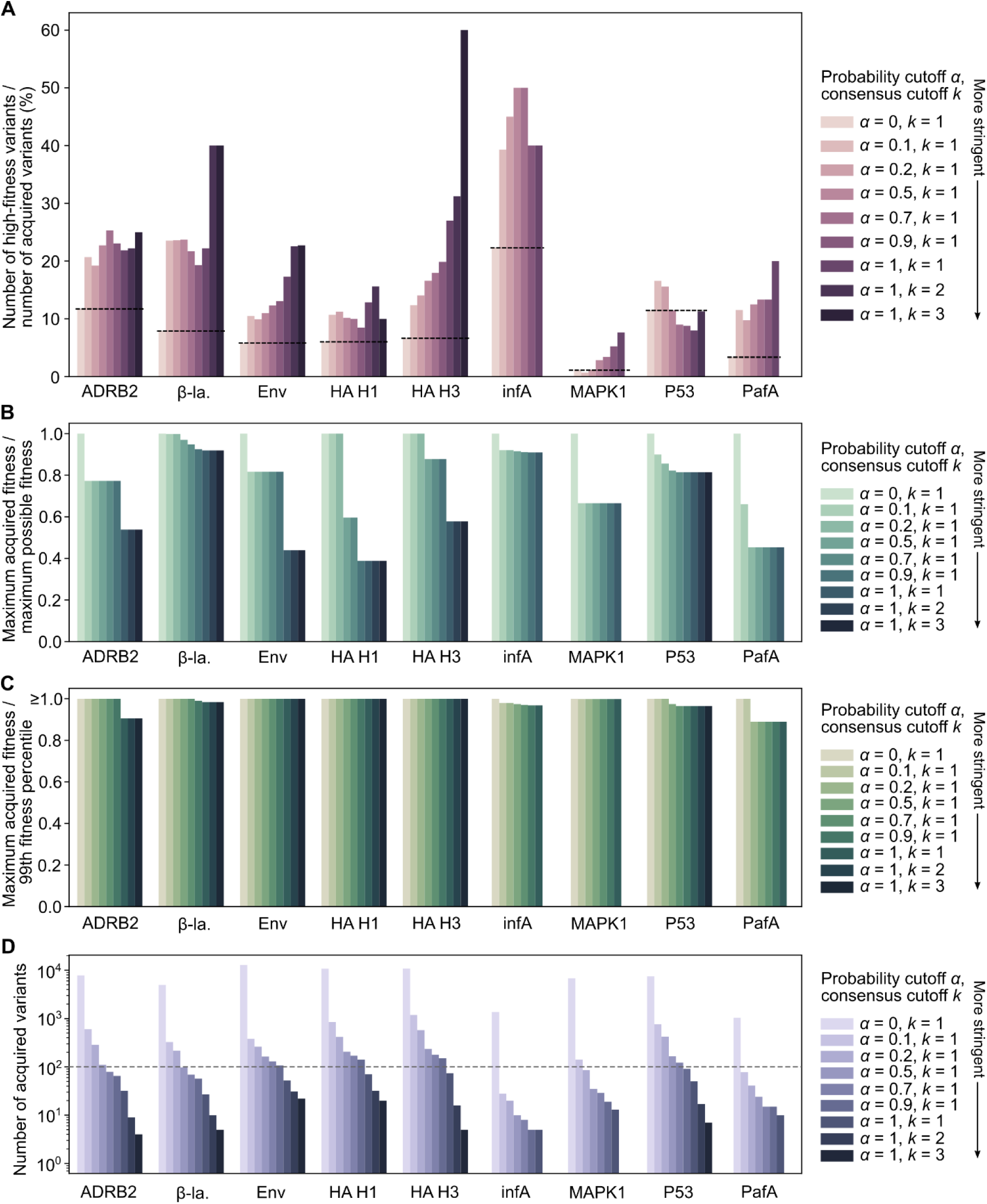
Relationship between likelihood stringency and extrinsic fitness efficiency. To obtain the set 𝒜 of language-model-recommended variants, we varied two parameters controlling the stringency of acquired variants (where more stringent corresponds to fewer variants): *α* is a cutoff controlling the likelihood ratio of the mutant probability to the wildtype probability, and *k* is a cutoff controlling the number of consensus language models (**Methods**). (**A**) At varying cutoffs, we computed the percentage fraction of variants in 𝒜 that correspond to high-fitness variants, using scanning mutagenesis data for validation. When *α* = 0 and *k* = 1, this value is equivalent to the percentage of high-fitness variants in the full scanning mutagenesis dataset (a black dashed line is also drawn at this value for each protein). In all cases except for P53, we observe that increasing the likelihood stringency generally improves the efficiency at which high-fitness variants are acquired. In Figure 4, we report values for *α* = 1, *k* = 2, except for when these cutoffs result in |𝒜| < 5 (infA, MAPK1, and PafA), in which case we report *α* = 1, *k* = 1. (**B, C**) Given a set of acquired variants 𝒜 at varying cutoffs, we also computed how much the maximum fitness represented in 𝒜 compares either to the maximum possible fitness value obtained across the full mutational scan (**B**) or to the 99^th^ percentile of fitness values across the full mutational scan (**C**). To compare across proteins, we plotted the maximum acquired fitness value normalized by the maximum possible fitness (**B**) or by the 99^th^ percentile with a threshold at 1 (**C**). At even at the most stringent cutoffs, the best acquired variant of most proteins has at least 50% of the fitness value of the maximum fitness peak. Additionally, at the most stringent cutoffs, the best acquired variant of all proteins is above or close to the 99^th^ percentile of fitness values. (**D**) We plotted the number of acquired variants |𝒜|, which is the denominator of the values plotted in (**A**). A gray horizontal dashed line is also plotted at 100.

## Supplementary Information

### Antibody sequences

Below are the antibody protein sequences defined as wildtype in this study:

- MEDI8852 VH: QVQLQQSGPGLVKPSQTLSLTCAISGDSVSSYNAVWNWIRQSPSRGLEWLGRTYYRSGWYNDYAESVKSRITINPDTSKNQFSLQLNSVTPEDTAVYYCARSGHITVFG VNVDAFDMWGQGTMVTVSS
- MEDI8852 VL: DIQMTQSPSSLSASVGDRVTITCRTSQSLSSYTHWYQQKPGKAPKLLIYAASSRGSGVPSRFSGSGSGTDFTLTISSLQPEDFATYYCQQSRTFGQGTKVEIK
- MEDI8852 UCA VH: QVQLQQSGPGLVKPSQTLSLTCAISGDSVSSNSAAWNWIRQSPSRGLEWLGRTYYRSKWYNDYAVSVKSRITINPDTSKNQFSLQLNSVTPEDTAVYYCARGGHITIFG VNIDAFDIWGQGTMVTVSS
- MEDI8852 UCA VL: DIQMTQSPSSLSASVGDRVTITCRASQSISSYLNWYQQKPGKAPKLLIYAASSLQSGVPSRFSGSGSGTDFTLTISSLQPEDFATYYCQQSRTFGQGTKVEIK
- mAb114 VH: EVQLVESGGGLIQPGGSLRLSCAASGFALRMYDMHWVRQTIDKRLEWVSAVGPSGDTYYADSVKGRFAVSRENAKNSLSLQMNSLTAGDTAIYYCVRSDRGVAGLF DSWGQGILVTVSS
- mAb114 VL: DIQMTQSPSSLSASVGDRITITCRASQAFDNYVAWYQQRPGKVPKLLISAASALHAGVPSRFSGSGSGTHFTLTISSLQPEDVATYYCQNYNSAPLTFGGGTKVEIK
- mAb114 UCA VH: EVQLVESGGGLVQPGGSLRLSCAASGFTFSSYDMHWVRQATGKGLEWVSAIGTAGDTYYPGSVKGRFTISRENAKNSLYLQMNSLRAGDTAVYYCVRSDRGVAGLF DSWGQGTLVTVSS
- mAb114 UCA VL: DIQMTQSPSSLSASVGDRVTITCRASQGISNYLAWYQQKPGKVPKLLIYAASTLQSGVPSRFSGSGSGTDFTLTISSLQPEDVATYYCQKYNSAPLTFGGGTKVEIK
- S309 VH: QVQLVQSGAEVKKPGASVKVSCKASGYPFTSYGISWVRQAPGQGLEWMGWISTYNGNTNYAQKFQGRVTMTTDTSTTTGYMELRRLRSDDTAVYYCARDYTRGAW FGESLIGGFDNWGQGTLVTVSS
- S309 VL: EIVLTQSPGTLSLSPGERATLSCRASQTVSSTSLAWYQQKPGQAPRLLIYGASSRATGIPDRFSGSGSGTDFTLTISRLEPEDFAVYYCQQHDTSLTFGGGTKVEIK
- REGN10987 VH: QVQLVESGGGVVQPGRSLRLSCAASGFTFSNYAMYWVRQAPGKGLEWVAVISYDGSNKYYADSVKGRFTISRDNSKNTLYLQMNSLRTEDTAVYYCASGSDYGDYL LVYWGQGTLVTVSS
- REGN10987 VL: QSALTQPASVSGSPGQSITISCTGTSSDVGGYNYVSWYQQHPGKAPKLMIYDVSKRPSGVSNRFSGSKSGNTASLTISGLQSEDEADYYCNSLTSISTWVFGGGTKLTVL
- C143 VH: EVQLVESGGGLVQPGGSLRLSCAASGFSVSTKYMTWVRQAPGKGLEWVSVLYSGGSDYYADSVKGRFTISRDNSKNALYLQMNSLRVEDTGVYYCARDSSEVRDHP GHPGRSVGAFDIWGQGTMVTVSS
- C143 VL: QSALTQPASVSGSPGQSITISCTGTSNDVGSYTLVSWYQQYPGKAPKLLIFEGTKRSSGISNRFSGSKSGNTASLTISGLQGEDEADYYCCSYAGASTFVFGGGTKLTVL

